# The Role of Conformational Changes in TcmN Aromatase/Cyclase in Polyketide Biosynthesis

**DOI:** 10.64898/2026.02.27.708631

**Authors:** Veronica S. Valadares, Ana Clara Granja, Luan C. Martins, Krishna Padmanabha Das, Elio A. Cino, Mariana T. Q. de Magalhães, Ana Paula Valente, Haribabu Arthanari, Adolfo H. Moraes

## Abstract

Polyketide biosynthesis relies on the conformational adaptability of type II polyketide synthases and accessory enzymes, which direct chain folding and regiospecific cyclization. The aromatase/cyclase TcmN from *Streptomyces glaucescensis* catalyzes the first two ring closures of tetracenomycin C. Still, the molecular basis by which conformational dynamics regulate substrate binding and product release remains unresolved. Understanding how conformational transitions control ligand recognition and prevent aggregation is crucial for deciphering the molecular bases of polyketide biosynthesis and for guiding engineering strategies to synthesize novel natural products. Here, we investigated how ligand interactions modulate the conformational equilibrium of TcmN and the mechanistic consequences for catalysis. Using NMR spectroscopy (STD, CSP, relaxation dispersion), calorimetry, molecular docking, and microsecond-scale molecular dynamics simulations, we mapped the conformational ensembles of apo TcmN and its complexes with naringenin (a substrate/product analogue) and intermediate 12 (INT12). Apo TcmN samples both open and closed conformations. Naringenin preferentially stabilizes the closed state, a conformation thought to protect hydrophobic residues from solvent exposure. In contrast, INT12 shifts the equilibrium toward the open state, characterized by an expanded cavity that permits substrate entry, product release, and accommodation of extended intermediates. Hydrogen-bond analysis highlighted conserved catalytic residues (R82, E34, Q110, T133) as key anchors for productive poses. These results establish that TcmN functions through a ligand-gated breathing mechanism, in which successive intermediates selectively tune the cavity volume and shape, balancing catalytic efficiency with protection against aggregation. Conformational adaptability emerges as a central determinant of aromatase/cyclase function, providing molecular insights relevant for polyketide biosynthetic engineering.

**Graphical Abstract:** 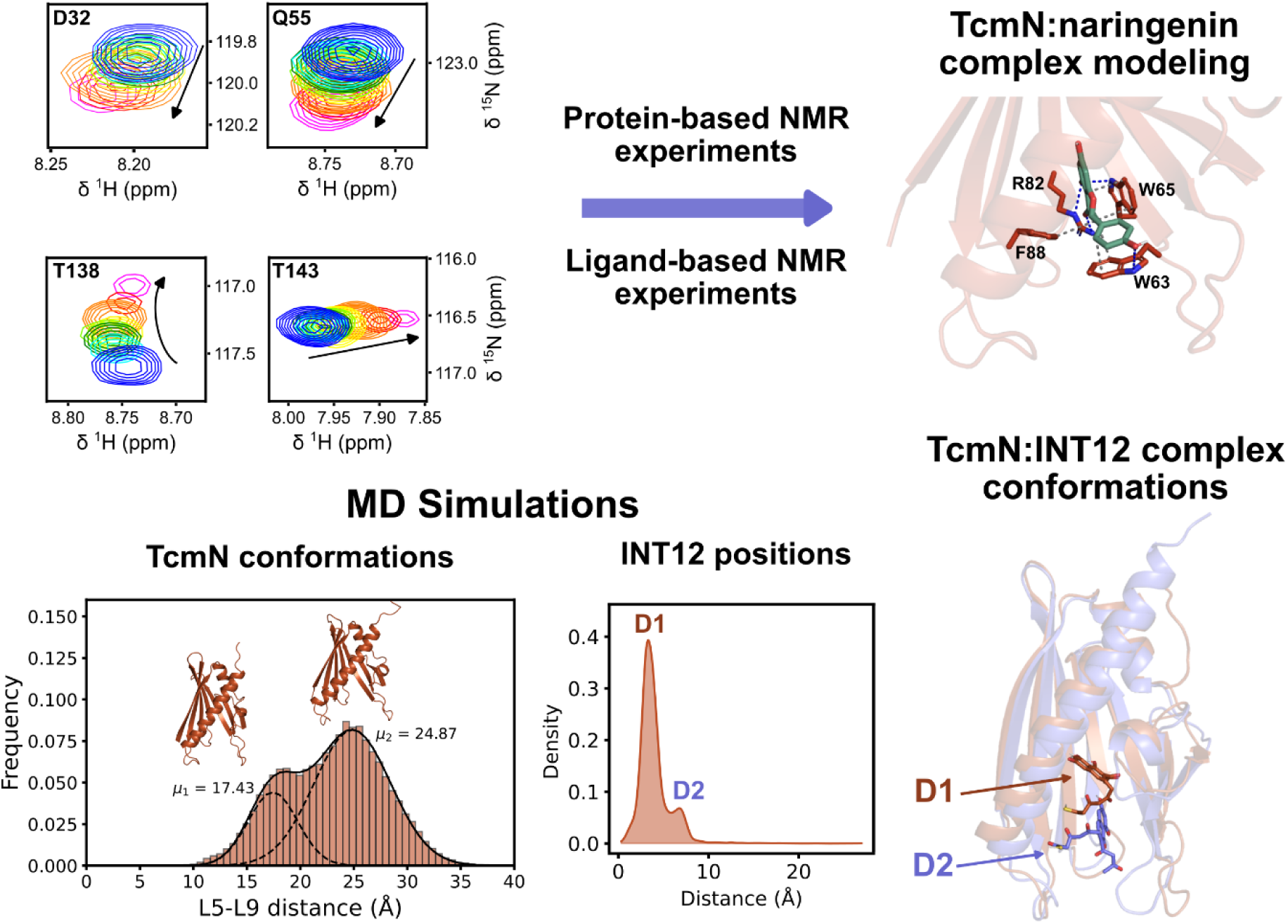

## INTRODUCTION

Polyketides are a diverse group of natural products with various chemical structures and pharmaceutical applications, produced by bacteria, fungi, and plants^1^. Polyketide synthases (PKS) catalyze the formation of polyketides by condensing acetyl coenzyme A via malonyl-CoA. Type II PKSs are responsible for producing aromatic polyketides, and are composed of separate mono and bi-functional enzymes^2^. TcmN, an enzyme from *Streptomyces glaucescensis*, plays a crucial role in the cyclization, aromatization, and folding of polyketides by catalyzing the formation of the first and second rings (Fig. 1A and B) ^3,4^. The minimal functional composition of type II PKSs, known as MinPKS, includes a ketosynthase, a chain length factor, an acyl carrier protein (ACP), and a malonyl transferase. It produces a highly reactive linear poly-β-keto intermediate. Specific components of type II PKS promote its correct aromatization and folding. In a non-reducing system, the growing poly-β-ketone intermediate is transported directly from the ketosynthase to the aromatase/cyclase (ARO/CYC) by ACPs. MinPKS from different organisms can be combined with TcmN to produce distinct polyketides, making the system suitable for creating novel compounds^5–8^.

**Figure 1.**
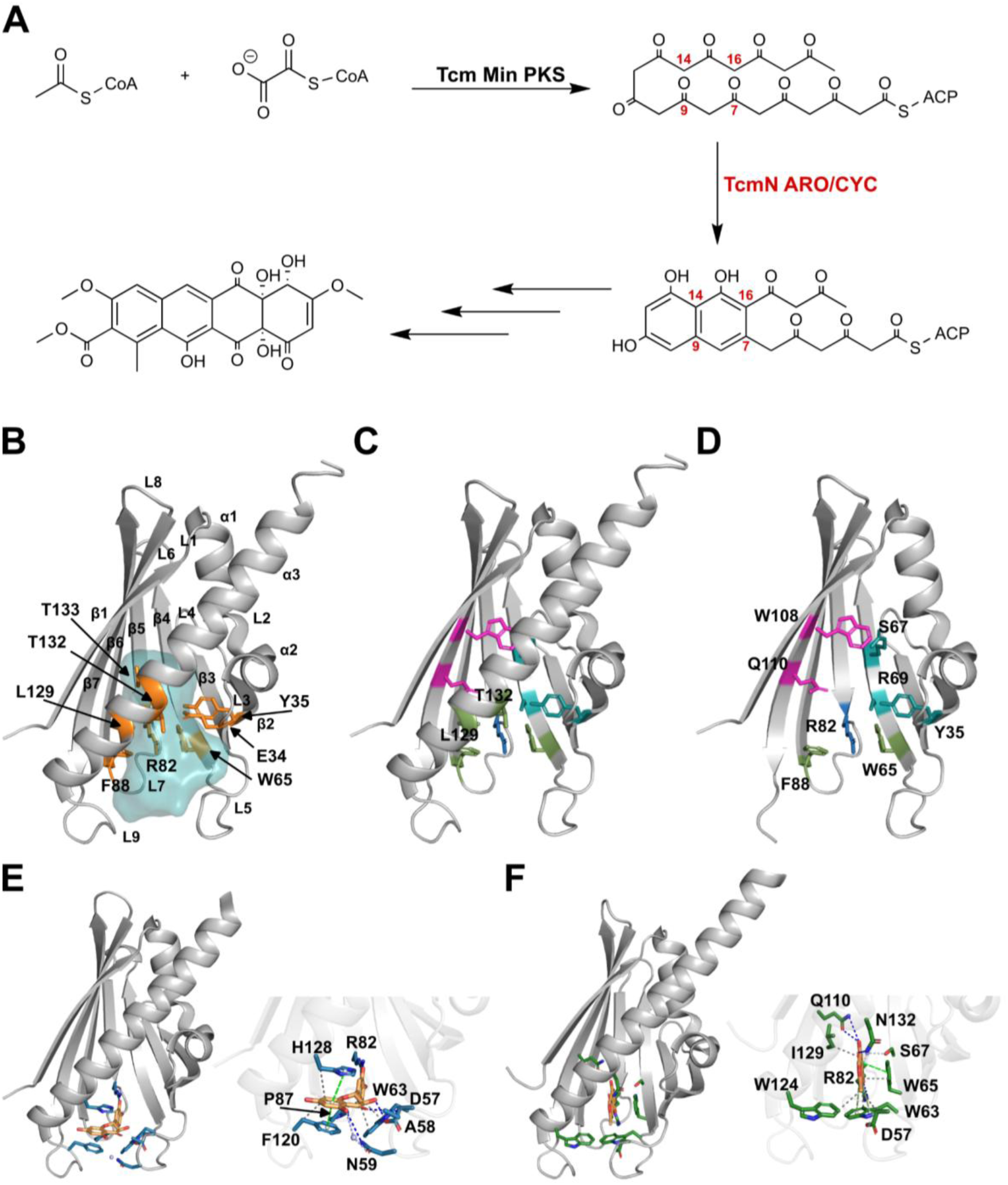
Structural features of Bet v 1–like aromatase/cyclases. (A) Simplified biosynthetic pathway of tetracenomycin C. Adapted from Ames, B. et al., Proceedings of the National Academy of Sciences April 2008, 105 (14) 5349-5354; © 2008 by The National Academy of Sciences of the USA. (B) Crystal structure of TcmN (PDB ID: 2RER), highlighting secondary structure elements, the main cavity volume (light blue), and key residues at the cavity bottleneck (orange). (C-D) Crystal structure of TcmN Aro/Cyc with residues critical for the catalytic colored in magenta (polyketide folding), green (cavity neck), cyan (first cyclization), and blue (second cyclization). (E) Crystal structure of TcmN Aro/Cyc in complex with trans-dihydroquercetin (taxifolin) (PDB ID: 3TVQ), with interacting side chains shown in blue and taxifolin in orange. (F) Crystal structure of WhiE ORFVI Aro/Cyc in complex with JRO (PDB ID: 3TL1), with interacting side chains in blue and JRO in orange. Interactions of TcmN with taxifolin and WhiE ORFVI with JRO were found using the PLIP web server (https://plip-tool.biotec.tu-dresden.de/plip-web/plip/index).

TcmN has a β-sandwich fold, with a long C-terminal α-helix and two small helices between β1 and β2 forming a helix–loop–helix motif (Fig. 1B, C, and D). Nascent polyketide intermediates bind to the interior pocket of TcmN for regiospecific cyclization and aromatization. The cavity has a ∼20 Å deep interior with a distribution of hydrophobic and polar residues that can accommodate an ACP-bound decaketide (20 carbons)^3,4^. The cavity has a narrow neck defined by residues W65, F88, L129, Y35, T132, T133, E34, and R82. Pocket residues W28, F32, W65, S67, R69, M91, and W95 are highly conserved among ARO/CYCs, while T54, L93, H128, T132, T133, and N136 are unique to TcmN. R69, T35, W108, and Q110 orient the polyketide chain inside the cavity, while S67 and R82 anchor the polyketide (Fig. 1C and D). The size and shape of the TcmN cavity modulates chain folding for C9-C14 first-ring cyclization. C9-C14 first-ring cyclization and interactions with residues in the narrow neck region orient C7 and C16 so that the C7 carbonyl hydrogen bonds with R82. The C16 methylene of the polyketide is proposed to position near TcmN E34 and Y35, favoring C7-C16 second-ring cyclization. The TcmN cavity might accommodate up to three linearly fused rings. Third-ring cyclization likely occurs spontaneously. Afterward, the polyketide is transferred to TcmI, a cyclase that is structurally and functionally distinct from TcmN, where fourth-ring cyclization occurs to produce a compound that undergoes further enzymatic modification to yield the anthracycline antibiotic Tetracenomycin^3,6^.

Functional insights were provided by site-directed mutagenesis. Ames et al.^3^ demonstrated that residues R69, Y35, R82, W108, and Q110 are essential for catalysis, with Q110 positioned at the cavity bottleneck, where its side chain mediates hydrogen bonding with reaction intermediates. W95, located in the cavity interior, interacts with W108 through π-stacking, a contact proposed to contribute to the structural and conformational stability of the enzyme (Fig. 1C, D, and E). Other studies emphasized the importance of residues at the cavity entrance: F120 and W63, located in loops L5 and L9, were implicated in regulating the opening and closing motions of the cavity, which are required for substrate access and product release^4^.

Conformational changes play a central role in enzyme–substrate recognition, as demonstrated for other Bet v 1–like proteins^9–12^. For TcmN, transient sampling of open states appears essential for substrate binding and may be equally critical for catalytic activity. The TcmN cavity is lined with hydrophobic residues that, if excessively exposed to solvent, could promote aggregation. Valadares et al.^4^ proposed a model describing the conformational equilibrium of TcmN between a closed and an open state. In this model, the enzyme predominantly adopts a closed conformation, while transient opening enables substrate entry and product release. Such kinetic and thermodynamic fine-tuning not only facilitates efficient ligand exchange but also minimizes the lifetime of the aggregation-prone open state, thereby reducing the risk of misfolding and amorphous aggregation.

Structural studies have provided further insights into TcmN binding mechanisms. Lee et al.^6^ solved the cocrystal structure of TcmN with trans-dihydroquercetin (taxifolin), which was positioned near the entrance of the catalytic pocket rather than deeply buried inside. Taxifolin binding induced conformational rearrangements in residues R82, H128, and F120, as well as in loops L5 and L7. The binding mode involved aromatic interactions with W63, W65, F88, and F120; hydrophobic contacts with M125 and P87; and hydrogen bonds with R82 and D57 (Fig. 1E). The orientation of the first aromatic ring, pointing outward from the cavity, was suggested to reflect a possible product-release mode, in which R82 may function as a hydrogen-bond regulator of substrate positioning and product exit. Consistently, structural analyses of the homologous aromatase/cyclase WhiE-ORFVI revealed that binding of the polyketide mimic JRO induces significant conformational changes in loops 5, 7, and 9, and particularly in W63, which undergoes a ∼4 Å displacement and ∼50° ring rotation to establish perpendicular π–π stacking with the ligand^6^ (Fig. 1F). This structural evidence supports the proposal that W63 may function as a molecular gate controlling substrate entry or product exit^13^, a role likely conserved in TcmN, given the high sequence and structural similarity between the two enzymes.

Despite the advances in describing the open–closed conformational equilibrium of TcmN, an important question remains: how does this equilibrium regulate substrate binding, modulate the enzyme’s catalytic function during the successive steps of polyketide aromatization/cyclization, and ultimately control product release?

In this study, we investigate the conformational changes underlying TcmN molecular recognition using NMR spectroscopy to monitor interactions with the intermediate/product analog naringenin. These interactions were analyzed by saturation transfer difference (STD) and chemical shift perturbation (CSP) NMR experiments. Molecular docking of the TcmN–naringenin complex provided a framework for modeling the interaction with intermediate 12 (INT12), an intermediate of tetracenomycin C and a product of TcmN aromatization/cyclization catalysis, as suggested by Ames et al.^3^. Molecular dynamics simulations were then performed for apo TcmN, the TcmN–naringenin complex, and the TcmN–INT12 complex, allowing us to characterize the enzyme’s conformational plasticity. The results reveal that apo TcmN samples two major conformations: a closed and an open state. The binding of naringenin stabilizes the closed conformation, while INT12 shifts the equilibrium toward the open state and further promotes a variation of the closed state. Together, these findings demonstrate that the conformational adaptability of the Bet v 1-like fold of TcmN is essential for its catalytic function, as distinct cavity volumes and shapes are required for substrate recognition, cyclization, and aromatization steps.

## MATERIALS AND METHODS

### Protein expression and purification

The gene encoding N-terminally His_6_-tagged TcmN (residues 1–173) was commercially synthesized and cloned into a pET28a plasmid by GenScript (Piscataway, USA). *E. coli* BL21 (DE3) cells were transformed with the construct and initially cultivated at 37 °C in a 1:10 starter culture ratio of Luria–Bertani (LB) medium supplemented with 50 µg·mL^−1^ kanamycin for 16 h with shaking at 200 rpm. The culture was used to inoculate 500 mL of LB medium. For isotope labeling, M9 minimal medium containing ^15^N–ammonium chloride and ^13^C–glucose (Sigma-Aldrich, USA) was used to produce ^15^N- and ^15^N/^13^C-labeled samples. Protein expression was induced with 0.5 mM isopropyl-β-D-thiogalactopyranoside (IPTG) at an optical density of 0.8 (600 nm), and cultures were incubated for 16 h at 22 °C with shaking at 200 rpm. Cells were harvested by centrifugation at 4,500 rpm for 40 min at 4 °C using a Thermo Scientific TX-750 swinging-bucket rotor (Thermo Fisher, USA). The pellet was resuspended in 50 mM sodium phosphate buffer, 300 mM NaCl, 10 mM imidazole, and pH 8.0, lysed, and centrifuged at 11,000 rpm for 40 min at 4 °C.

The supernatant was subjected to Ni^2+^ affinity chromatography using a HisTrap HP Ni-NTA column (GE Healthcare, USA), which was equilibrated in 50 mM sodium phosphate buffer, 300 mM NaCl, 40 mM imidazole, and pH 8.0. Ni^2+^-Bound protein was eluted with an imidazole gradient up to 500 mM. The His_6_-tag was cleaved with thrombin according to the manufacturer’s protocol (Thrombin CleanCleave Kit, Sigma-Aldrich, USA). The protein was subsequently dialyzed against a 20 mM sodium phosphate buffer, pH 7.0, and 150 mM NaCl. Typical yields were 22 mg purified TcmN per liter of culture, with>95% purity. Protein analysis was carried out by SDS–PAGE, and protein concentration was determined by UV–Vis absorbance at 280 nm, using an extinction coefficient of 37,470 M^−1^·cm^−1^, calculated from the TcmN primary sequence with ProtParam (https://web.expasy.org/protparam/).

### TcmN-naringenin interaction by differential scanning calorimetry (DSC)

Differential scanning calorimetry (DSC) experiments were performed on a MicroCal PEAQ DSC instrument (Malvern, UK) at the *Laboratório Multiusuário do Departamento de Biofísica* of the Federal University of São Paulo. Thermograms were acquired using 211 μM TcmN in a 20 mM sodium phosphate buffer, pH 7.0, 150 mM NaCl. The thermal stability of naringenin-bound TcmN was evaluated by recording DSC thermograms (replicates, n = 3) in the presence of 1 or 2 mM naringenin. Since the naringenin stock solution was prepared in DMSO, final protein samples contained 4% or 8% (v/v) DMSO, respectively.

The effect of naringenin binding on thermal resistance was performed in triplicate at a controlled scan rate of 1.0 °C·min^−1^ using a 300 μL capillary cell. The reference cell was filled with a buffer containing the same concentrations of naringenin and DMSO as in the measurement cell. Reversibility of thermal transitions was assessed by reheating the samples immediately after cooling. DSC data were analyzed using Origin 2018 (OriginLab Corporation, USA). Protein thermal stability was described by the temperature of the maximum transition (*T_m_*).

### Isothermal titration calorimetry (ITC) measurements

ITC experiments were conducted at 25 °C using an Affinity ITC instrument equipped with an autosampler (TA Instruments, New Castle, DE). Measurements were performed in a 20 mM sodium phosphate buffer (pH 7.0) containing 150 mM NaCl and 4% (v/v) DMSO. A 140 μM protein solution (or buffer for control experiments) was placed in the calorimetric cell and titrated with 2.5 μL injections of 2 mM ligand solution at 350 s intervals, with stirring at 125 rpm. The resulting isotherms were corrected by subtracting buffer–ligand control titrations and fitted to a single-site binding model using NanoAnalyze software (TA Instruments) to determine the dissociation constant (*K_d_*).

### NMR spectroscopy

#### Protein backbone assignment

Protein backbone NMR assignment experiments were performed at 298 K using Bruker 900 MHz Avance Neo and 800 MHz Avance III spectrometers equipped with TXI 5 mm triple-resonance probe and a Bruker Avance Neo 600 MHz spectrometer equipped with a PABBO 5 mm SmartProbe. For the TcmN backbone resonance assignment, the following NMR triple-resonance experiments were acquired and analyzed: ^1^H-^15^N HSQC, ^1^H-^13^C HSQC, HNCO, HN(CA)CO, HNCA, HNCACB, and CBCA(CO)NH. 2D experiments were acquired using the traditional full acquisition schedule, and 3D experiments using non-uniform Poisson Gap Sampling^14,15^. The spectra were processed using Topspin 3.7 (Bruker Corporation, USA) and NMRpipe^16^ and analyzed using CCPN analysis 2.4.2^17^.

#### NMR relaxation experiments

The backbone dynamics of TcmN were monitored using longitudinal (R1) and transverse (R2) relaxation rates on a Bruker Avance Neo 600 MHz spectrometer. The overall protein correlation time, τ_c_, was calculated from the *R_2_/R_1_* ratio using the following equation:

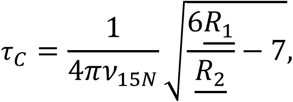

where 𝜈_15𝑁_ is the ^15^N Larmor frequency, ̲𝑅_1_ and ̲𝑅_2_ are the average relaxation rates obtained from 1D version ^15^N R_1_ and R_2_ NMR relaxation experiments.

Dynamics on the *μs-ms* timescale were monitored by NMR relaxation-compensated ^15^N single-quantum relaxation compensated Carr-Purcell-Meiboom-Gill relaxation dispersion experiments (^15^N-RD-CPMG)^18^. ^15^N-RD-CPMG experiments were acquired at 298 K using a Bruker Avance Neo 600 MHz spectrometer, equipped with a 5 mm dual-channel multinuclear Smart Probe, located at the Nuclear Magnetic Resonance Laboratory (LAREMAR) of the Federal University of Minas Gerais. Experimental *R_2_* effective rates, *R_2,eff_ (s^-^*^1^), were obtained from ^1^H-^15^N correlation spectra intensities using the software CCPN Analysis 2.4.2^17^, and applying the following equation:

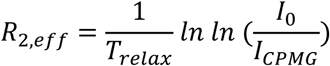

where *T_relax_* is the relaxation delay of 30 ms, *I_CPMG_* is the intensity measured in ^1^H-^15^N spectra acquired with CPMG frequency, ν_CPMG_, ranging from 66.7 to 1000 Hz. *I_0_* is the peak intensity obtained with T_relax_ of 0 ms. Experimental errors were calculated from the standard deviation of the noise in the resonance intensities, following Loria et al. ^18^.

#### Interaction of TcmN with Naringenin and Taxifolin by NMR spectroscopy

The saturation transfer difference (STD) NMR experiment was employed to evaluate the interaction between TcmN and the ligands. One-dimensional ^1^H STD spectra were recorded at 25 °C. Samples (600 μL) contained 20 μM TcmN in 20 mM sodium phosphate buffer (pH 7.0), 150 mM NaCl, 1 mM naringenin or taxifolin, 4% (v/v) DMSO-d_6_, and 10% (v/v) D_2_O, corresponding to a protein-to-ligand molar ratio of 1:20.

STD build-up curves were acquired for TcmN-naringenin interaction using saturation times of 0.25, 0.75, 1.5, 2.0, 2.5, 3.0, 3.5, and 4.0 s. Selective saturation was applied using a train of 50-ms Gaussian pulses^19^. The on- and off-resonance irradiation frequencies were set to 0 ppm and −40 ppm, respectively. STD signal intensities obtained at different saturation times were used to calculate STD amplification factors^20^ according to the following equation:

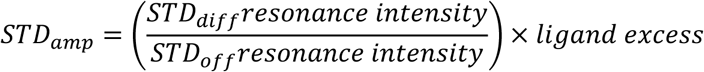

The interaction of TcmN with naringenin was monitored by NMR titration experiments, in which the ligand was incrementally added to isotopically ^15^N-labeled protein samples. For each titration point, a ^1^H–^15^N HSQC spectrum was acquired using the hsqcetfpf3gp pulse sequence. Experiments were conducted at 298 K on a Bruker Avance III 800 MHz spectrometer equipped with a TXO cryogenic probe, located at the Harvard Medical School Biomolecular NMR Facility and the Dana-Farber Cancer Institute NMR Core, and on a Bruker Avance Neo 600 MHz spectrometer equipped a 5 mm dual-channel multinuclear Smart Probe, located at the Nuclear Magnetic Resonance Laboratory (LAREMAR) of the Federal University of Minas Gerais.

NMR Samples were prepared in 20 mM sodium phosphate buffer, pH 7.0, 150 mM NaCl, and 10% (v/v) D2O. At each titration point, TcmN was mixed with a solution of naringenin dissolved in DMSO-d_6_. The final protein and ligand concentrations used are summarized in Tables S1 and S2.

The chemical shift perturbation (CSP) index was calculated using the following equation:

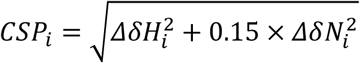

where *ΔδH_i_* and *ΔδN_i_* are the chemical shift variation in the ^1^H and ^15^N dimension, respectively.

#### Line shape analysis of the TcmN NMR titration experiments

The NMR signal shape analysis was performed using the software TITAN^21^. They were implemented in MATLAB. The analyses were conducted according to the methodology described by Waudby et al^22^.

For TITAN analysis, signals of interest were selected by manually defining polygonal regions of interest (ROIs) that encompassed the entire trajectory of each peak across the titration series, as observed in the overlaid spectra. Within each ROI, the chemical shift positions of the free and bound states were manually assigned. Only resonances from residues exhibiting significant chemical shift perturbations (CSPs), higher than the mean plus one standard deviation, and free of peak overlap were retained for line shape analysis.

Given the large number of variables, data fitting was performed in two steps. In the first step, only the chemical shift and linewidth of the free protein resonance were optimized, while kinetic constants were kept fixed. Default initial and maximum values from TITAN were used for the kinetic parameters, and the *K_D_* was fixed at 140 µM, as determined by ITC. In the second stage, chemical shifts, linewidths, and both kinetic and thermodynamic parameters were refined, except for the free protein chemical shift, using all spectra from the titration series. This procedure was applied to each selected resonance individually.

Thermodynamic and kinetic parameters of the interaction model were obtained from global fitting and spectral simulation of the selected signals. Model suitability and fit quality were assessed by comparing TITAN simulations with experimental spectra. Parameter uncertainties were estimated using a bootstrap resampling procedure with 100 replicas. Four interaction models (lock-and-key, conformational selection, induced fit, and sequential binding) were evaluated for their ability to reproduce the experimental NMR signal behavior observed during the titration of naringenin^21,22^.

### Molecular Docking simulations

To select poses for seeding MD simulations to study TcmN and interactions with intermediate and product analogs, docking simulations were performed using High Ambiguity Driven biomolecular DOCKing (HADDOCK) version 2.4 and AutoDock Vina 1.2^23^. The TcmN crystal structure (PDB ID: 2RER) was selected for docking simulations. The chemical structures of (*S*)- and (*R*)-naringenin were modeled using MarvinSketch ^19.21^ software. The INT12 structure without ACP was modeled using MarvinSketch ^19.21^. TcmN residues E34, Y35, S67, R89, W95, H128, and T133 (predicted by Ames et al. 2008 as pocket residues that make contact with ligands) and residues A80, M91, W108, Q110, I140, T132, L129 and F112 (mapped by CSP experiment with titration of Naringenin to TcmN) were selected as HADDOCK restraints and defined as the protein active and passive residues, respectively, in the rigid-body docking step (it0). The mentioned residues were set as passive in the semi-flexible docking step (it1) and explicit solvent refinement step (water). 10000 poses were generated in it0 and 1000 poses in it1. The best-docked complexes of top-ranked clusters were selected and visualized using Discovery Studio Visualizer (BIOVIA, Dassault Systèmes, San Diego, USA, 2022) and UCSF Chimera^24^. Ligand poses from Haddock were aligned using GROMACS tools (gmx trjconv) using TcmN Cα for alignment. Pairwise RMSD, RMSD to the best-scoring pose, and ligand center-of-mass were calculated with GROMACS tools. The pairwise RMSD matrix for each ligand was embedded into a bi-dimensional space using T-distributed Stochastic Neighbor Embedding (t-SNE)^25^. Bidimensional embedding was then visually inspected and clustered using Density-based spatial clustering of applications with noise (DBSCAN)^26^, with ε set to 13 and a minimum of 75 samples to define a cluster. Cluster elements were visually inspected and selected based on their complementarity to the binding pocket, any polar interactions, and the pose score. For TcmN-INT12 interaction modelling, a semi-flexible docking simulation was also performed using the software Autodock Vina 1.2, positioning the center of the docking box at the center of mass of residue R82, at the center of the TcmN cavity, and a key residue for the interaction of TcmN with substrates, intermediates, and products. To maximize ligand sampling, AutoDock Vina 1.2 docking was performed with exhaustiveness set to 128 and up to 20 poses per run.

### Molecular Dynamics simulations

Ten independent replicates of 1 µs MD simulations were performed for free TcmN, Naringenin-bound TcmN, and INT12-bound TcmN using the software GROMACS 5^27^, employing the CHARMM36m^28^ force field and the standard TIP3P water model^29^. The TcmN crystal structure (PDB ID: 2RER) was used with protonation states of titratable residues predicted by PropKa^30^ at pH 7. Na^+^ and Cl^-^ were added to achieve charge neutrality and simulate a physiological concentration of 0.15 M.

Following steepest-descent minimization, 10 replicates were initiated with different velocity distributions, equilibrated and propagated using the same, previously described protocol^31^. First, initial atomic velocities were generated from a Maxwell-Boltzmann distribution at 298 K. Long-range electrostatic interactions were evaluated using the Particle Mesh Ewald method with a 1.2 nm real-space cutoff. Cutoff for van der Waals interactions was 1.2 nm, with a force-switch from 1.0 to 1.2 nm. All hydrogen-containing bonds were constrained using the LINCS algorithm. An initial 125 ps equilibration was performed in the NVT ensemble at 298 K using the Nose-Hoover thermostat while applying position restraints to the solute backbone (400.0 kJ/mol*nm) and side chains (40.0 kJ/mol*nm). Then, 125 ps unrestrained NPT equilibration was run using Parrinello-Rahman barostat with a 1.0 bar pressure. Then, the systems were propagated for 1 µs each in the NPT using the Parrinello-Rahman barostat (1.0 bar) and the Nose-Hoover thermostat (298 K).

For the TcmN complexed with naringenin and INT12, three poses obtained by docking simulation were selected as described above and used as the starting structures for the MD simulations. GROMACS analysis tools and in-house scripts were employed to analyze the trajectories. Principal component analysis (PCA) was performed using Python and the scikit-learn library on the pooled frames from the ten replicas, with least-squares fitting of the Cα atoms to the initial structure^32^. Due to high flexibility, residues K141 to P155 from the C-terminal were not considered in the PCA analysis. Representative structures of the closed and open states were taken from the centermost frames of the respective energy minima. Bidimensional histograms were prepared by binning the trajectory projections on the first two principal components and expressed in energy units using Boltzmann’s factor^33^. CASTp 3.0^34^ and 3V^35^ were used to identify and characterize cavities. The distance between loops L5 and L9 was evaluated by measuring the distance between the Cα of P119 (L9) and R61 (L5). Interpolation of structures to represent the motion along PC1 was performed using gmx tools^27^. One hundred interpolated structures were generated and analyzed. Gaussian fitting to the histograms of the L5-L9 distances was performed using non-linear regression using Python 3 scripts.

## RESULTS AND DISCUSSIONS

### Naringenin binding mode in the TcmN cavity

TcmN promotes C9-C14 first-ring, C7-C16 second-ring cyclization, and aromatization of unreduced linear polyketides. The poly-β-keto intermediate is highly reactive, making it difficult to study experimentally with the enzyme TcmN. To overcome this problem, we investigated the interaction of TcmN with naringenin and taxifolin, which contain two rings and hydroxyl and carbonyl groups, resembling the structure of the cyclic polyketide intermediate/product in TcmN-catalyzed aromatization and cyclization reactions. Using ITC, we observed a dissociation constant of 140 μmol/L (Fig. S1). DSC experiments showed a 2°C increase in thermal resistance of TcmN upon titration with naringenin (Fig. S2).

The interaction of TcmN with naringenin and taxifolin was evaluated using ^1^H STD NMR experiments with the ligand candidates at a 20-fold excess over the TcmN concentration. The interaction between TcmN and taxifolin was weak, as indicated by low signal-to-noise STD signals (Fig. S3). For the naringenin, STD experiments revealed stronger STD signals for the aromatic protons H1 and H5 (Fig. S4), which were confirmed by the STD amplification build-up curves (Fig. 2A), indicating that the two fused rings of naringenin are positioned deeply in the TcmN cavity.

**Figure 2.**
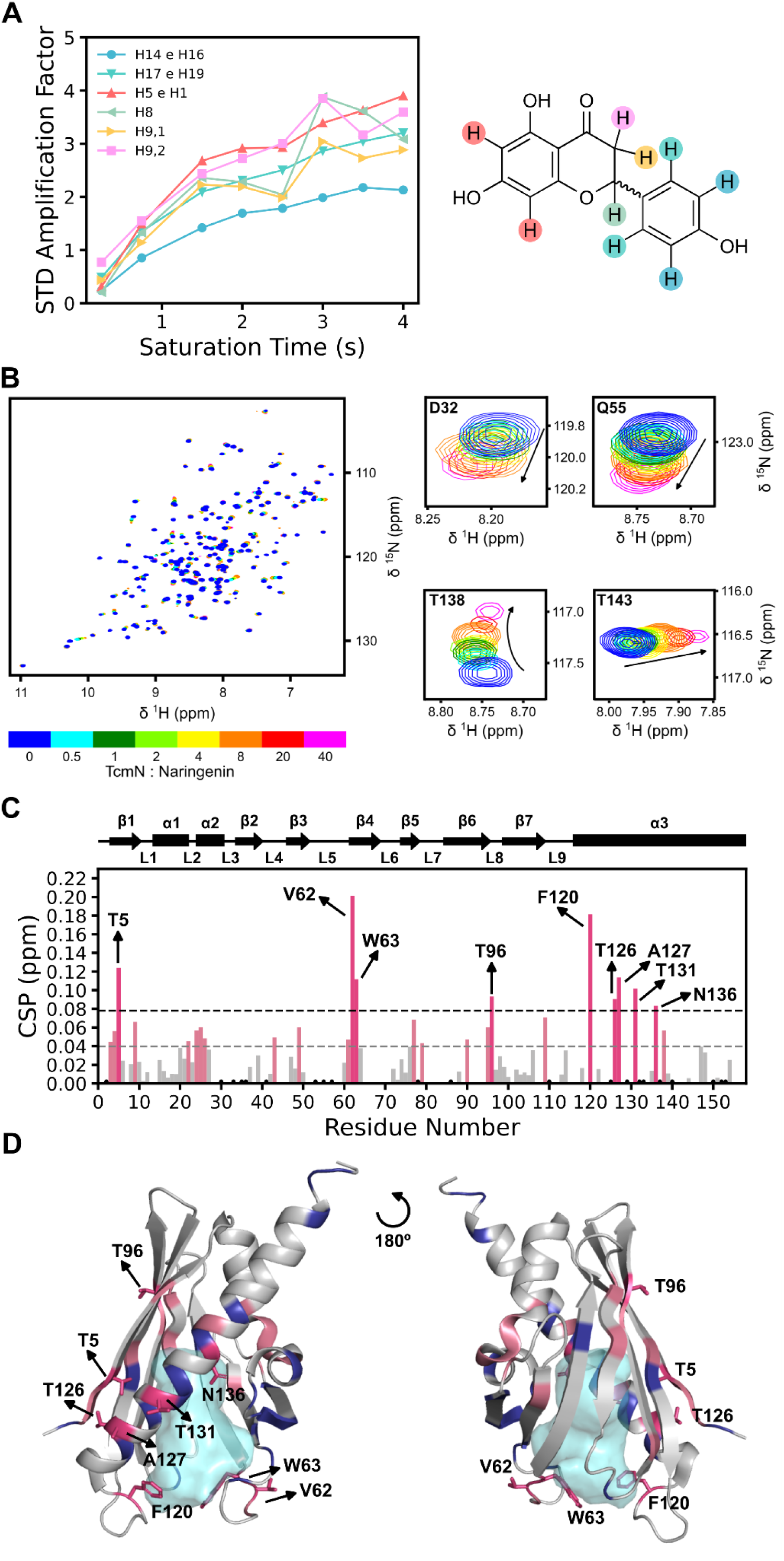
Naringenin binds to the central cavity of TcmN, as determined by NMR spectroscopy. (A) Saturation Transfer Difference (STD) NMR build-up curves confirm direct binding of naringenin to TcmN. Proton assignments for naringenin are shown. (B) Overlay of ¹H-¹⁵N HSQC spectra of TcmN titrated with naringenin, showing specific chemical shift deviations. The NMR signal modifications for residues D32, Q55, T138, and T143 are zoomed in. The color scale represents the [TcmN]:[naringenin] molar concentration ratio. (C) CSPs as a function of TcmN residue numbers. Residues with significant CSPs (above the mean + standard deviation, black dashed line) are colored in dark pink; broadened/disappeared residues are identified as black dots. (D) The TcmN structure colored by CSP, revealing the naringenin binding site (dark pink: highest CSP; light pink: above mean CSP; dark blue: broadened/disappeared residues).

To identify the interaction site in the TcmN structure, ¹H-¹⁵N HSQC spectra of ¹⁵N-labeled TcmN titrated with naringenin were acquired, and structural changes were determined by mapping the CSPs. Figure 2B shows an overlay of ¹H-¹⁵N HSQC of free and bound TcmN with different concentrations of naringenin. Significant CSPs were observed in residues T5, S33, Y35; D57, W63, F120, T126, N136, and M137 that surround the TcmN cavity and alpha-helix 3. Residues F120 and W63, located near the cavity entrance in loops 5 and 9, showed significant CSP (Fig. 2C). These loops, identified as regulators of cavity opening and closing movements necessary for substrate binding and product release, are crucial for TcmN function^4^. It can be inferred that residue W63 acts as a potential gatekeeper, similar to its homologous residue in WhiE-ORFVI, suggesting its role in controlling ligand access to the binding site^6^. Interestingly, the NMR signals of residues Y35, L129, and T132, which form part of the bottleneck region (Fig. 1B), broaden upon naringenin titration, implying a conformational equilibrium on an intermediate chemical shift timescale, caused by the naringenin interaction and the variation in cavity volume and shape. This variation in cavity volume and shape was also observed in structurally similar proteins, such as Bet v1 and Fag s 1^9,10,12^ . Additionally, the curved signal trajectories observed in the titration experiments indicate a more complex interaction between TcmN and naringenin, suggesting either the binding of multiple ligands or a more intricate mechanism, such as conformational selection or induced fit upon ligand binding. (Fig. 2B).

To gain deeper insight into the interaction mechanism, 2D line-shape analysis was performed using TITAN software on the ¹H–¹⁵N HSQC titration data for TcmN with naringenin. The titrations covered naringenin concentrations ranging from 0 to 561.6 µM at 600 MHz and from 0 to 2 mM at 800 MHz. Residues were selected for fitting when their CSPs exceeded the average value plus one standard deviation. Based on this criterion, the residues included were: T5, V62, T96, Q110, F120, and A127 at 600 MHz, and T5, V62, W63, T96, F120, T126, A127, T131, and N136 at 800 MHz (Fig. S5).

At lower ligand-to-protein ratios, the NMR signal trajectories displayed linear behavior, consistent with a simple two-state binding equilibrium (P + L ⇌ PL). The global fit under these conditions yielded a *K_D_* of 111 ± 5 μM, indicating a moderate affinity between TcmN and naringenin. At higher ligand excess, however, deviations from linearity became evident, and several residues exhibited curved titration trajectories. This nonlinearity suggests that additional binding or conformational events occur at elevated naringenin concentrations, inconsistent with a single-step binding model. These observations prompted the evaluation of alternative kinetic schemes—such as conformational selection, induced fit, and sequential binding—to determine which mechanism best reproduces the experimental line-shape behavior.

When analyzed with the software TITAN, single-site mechanisms, including the two-state, conformational selection, and induced-fit models, were unable to reproduce the experimental line shapes and chemical-shift migration patterns. Instead, only a sequential binding scheme (P + L₁ ⇌ PL₁; PL₁ + L₂ ⇌ PL₁L₂), representing stepwise occupation of a weaker second site, provided a satisfactory reproduction of both the line-shape evolution and the CSP migration throughout naringenin titration (Fig. S5).

To separate the two binding events, the 800 MHz dataset also fit with the ligand concentration restricted to ≤ 200 μM ([L]:[P] ≤ 4.02) (Fig. S6). Within this window, the TcmN:Naringenin interaction behaved as a two-state system, indicating the presence of a high-affinity naringenin-binding site, with a *K_D_* of (2 ± 5) μM. In the complete sequential binding analysis, the first binding step remained in the low micromolar range (*K_D_* ≈ 7 ± 5 μM). At the same time, the second was substantially weaker, with an estimated affinity in the millimolar range. These values should be interpreted with caution: the dissociation constant of the second site is difficult to determine precisely and may influence the apparent *K_D_* assigned to the first site in global fits. Nonetheless, the analysis indicates that TcmN binds to Naringenin via two distinct binding events, which differ by approximately two orders of magnitude in affinity. The NMR signal patterns further support the involvement of a second site. Enhanced shifts at F120 and W63, together with line broadening at D57 and M125, are consistent with secondary binding side in the entrance of the main cavity.

It is informative to compare the two naringenin-binding sites of TcmN identified in this study with findings from previous work on TcmN and another Bet v 1–like aromatases/cyclases. Lee, Ames, and Tsai⁶ determined the crystal structure of TcmN in complex with the flavonoid taxifolin. The crystal structure shows that the flavonoid occupies a TcmN entrance-region site (Fig. 1E), causing rearrangements of R82, H128, and F120; forming π-stacking interactions with W63, W65, F88, and F120; hydrophobic contacts with M125 and P87; and hydrogen interactions with R82 and D57 (Fig. 1E). The orientation of the first aromatic ring of taxifolin, pointing outward from the cavity, was suggested to reflect a possible product-release mode, in which R82 may function as a hydrogen-bond regulator of substrate positioning and product exit. Assuming a conserved flavonoid binding scaffold, it is likely that at low [L]:[P] ratios, Naringerin binds within the core pocket, and at high [L]:[P] ratios, it may occupy the entrance-region site, in a position like taxifolin.

### Conformational dynamics of TcmN upon Naringenin interaction by NMR relaxation experiments

The dynamics of free and naringenin-bound TcmN were investigated by ^15^N NMR relaxation measurements. Relaxation rates (*R_1_* and *R_2_*) were used to calculate the rotational correlation time (*τ_c_*), which provides information on protein size, shape, and oligomeric state. The *τ_c_* value of free TcmN was slightly higher than that of the naringenin-bound protein, suggesting that ligand binding may promote a more compact conformation. A decrease in *τ_c_* is consistent with a reduction in the hydrodynamic radius, as a more compact structure undergoes faster rotational tumbling in solution. Nevertheless, these results should be interpreted with caution, since the *τ_c_* values of free and bound TcmN fall within the experimental confidence interval. Relaxation rates and calculated τc values for both states are summarized in Table 1.

**Table 1:** Relaxation rates and correlation times of TcmN free and bound to Naringenin.

| TcmN state | $R_1$ ($\text{s}^{-1}$) | $R_2$ ($\text{s}^{-1}$) | $\tau_c$ (ns) |
| --- | --- | --- | --- |
| Free | $1.01 \pm 0.03$ | $7.8 \pm 0.9$ | $8.2 \pm 0.7$ |
| Bound | $0.96 \pm 0.08$ | $6.8 \pm 0.6$ | $7.8 \pm 0.8$ |

^15^N CPMG relaxation dispersion NMR spectroscopy^36,37^ was used to characterize residues of TcmN undergoing conformational exchange. Effective transverse relaxation rates (*R_2,eff_* (s^−1^)) obtained from CPMG experiments at 298 K for free TcmN (Figure S7A) showed no significant differences between data acquired at CPMG frequencies of 50 and 1000 Hz. In contrast, bound TcmN exhibited increased *R_2,eff_* (s^−1^) values at 50 Hz that were attenuated at 1000 Hz, approaching those observed for the free protein (Figure S7B). While CSPs highlight residues directly involved in protein–ligand interactions, *ΔR_2_* reflects conformational exchange between two TcmN states that remain in equilibrium even at high naringenin concentrations. These two states likely correspond either to (i) a high-affinity complex with one naringenin bound versus a second, lower-affinity state in which an additional ligand transiently occupies the entrance region, or (ii) a singly bound state in equilibrium with a minor free-like population. The observed *ΔR_2_* values therefore report on exchange between these two dominant conformers rather than on a change between free and fully saturated protein.

Taken together, the STD, titration, and ^15^N CPMG relaxation dispersion experiments identified both the naringenin protons and the TcmN residues involved in the interaction, providing key structural information for modeling the TcmN–naringenin and TcmN–INT12 complexes.

### Modelling TcmN complex with Naringenin and INT12 by molecular docking and molecular dynamics simulations

Ames et al.^3^ investigated the role of TcmN pocket residues through docking simulations with putative substrates and products, showing that the substrate interacts with residues that facilitate the catalysis of the first- and second-ring cyclizations of a linear polyketide chain. The TcmN pocket can accommodate up to three linearly fused rings, but not a fourth; third-ring cyclization was proposed to occur spontaneously³. To further explore the molecular recognition mechanism of TcmN with substrates, intermediates, and products, we performed molecular docking and molecular dynamics (MD) simulations. Among the intermediates predicted by Ames et al.^3^, we selected intermediate 12 (INT12) (Fig. 1A), which contains two linearly fused aromatic rings. The TcmN intermediate/product analog, naringenin, was also used for computational analyses. Because NMR interaction experiments were performed with racemic naringenin, docking and MD simulations were carried out with both R- and S-isomers using the crystal structure of TcmN (PDB ID: 2RER).

Docking was performed using HADDOCK v2.4. The center of mass of the ligand in each pose was analyzed to assess docking quality and binding position. Most R-naringenin (NAR-R) poses, and all S-naringenin (NAR-S) poses, are localized within the TcmN cavity, whereas no INT12 poses were positioned inside the pocket. For NAR-R and NAR-S, the best-scoring poses were used as references to calculate ligand RMSDs across all docking results. All poses were then aligned in GROMACS and clustered. NAR-S clusters generally exhibited lower energy scores than those of NAR-R. The centroid pose from each cluster was visualized, and the best candidates were selected based on agreement with STD-NMR results and docking scores. From NAR-S, pose 461w (score = –15.87) was selected, while from NAR-R, poses 71w (score = 23.94) and 568w (score = –8.16) were chosen (Figure S8).

Because HADDOCK failed to place INT12 inside the TcmN catalytic cavity—likely due to the size and high flexibility of the molecule, which reduces the efficiency of HADDOCK’s algorithm— AutoDock Vina was used for INT12 docking. In this case, several poses were generated with the ligand correctly buried in the cavity, and the highest-scoring pose was selected. INT12 and (S)-Naringenin docking poses are shown in Figures 3A-D.

**Figure 3.**
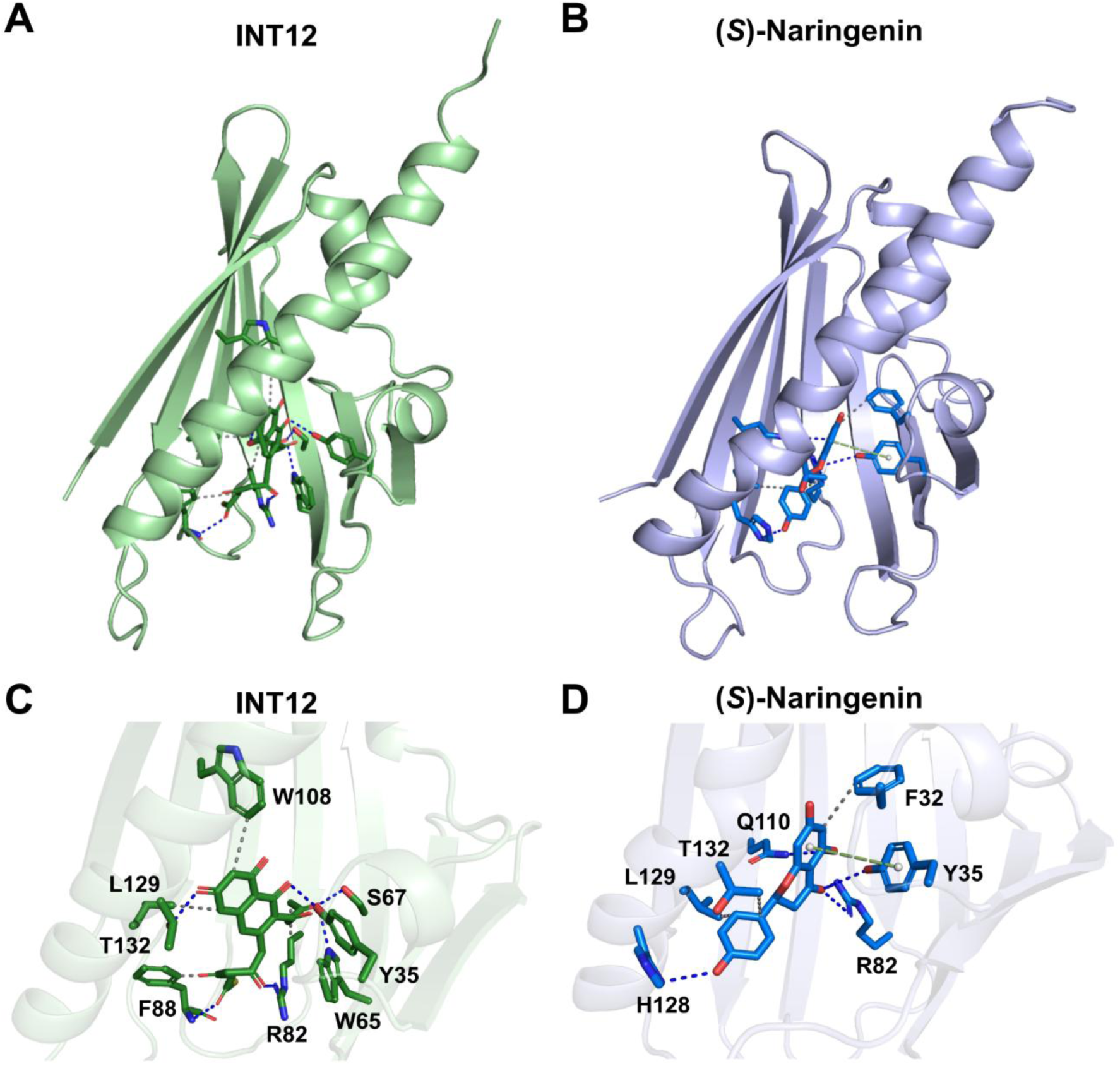
Molecular docking of INT12 and (S)-Naringenin. (A) Docking pose of INT12. (B) Docking pose of (S)-Naringenin. (C) Predicted INT12 interactions identified using the PLIP web server (https://plip-tool.biotec.tu-dresden.de/plip-web/plip/index). (D) Predicted (S)-Naringenin interactions identified using the PLIP web server.

Each selected pose of NAR-R, NAR-S, and INT12 bound to TcmN was used to initiate ten independent 1-µs MD simulations. In parallel, ten replicates of 1-µs MD simulations of apo TcmN were performed with the same force field to allow direct comparison of results. Trajectories from all replicates were analyzed, including free TcmN, INT12-bound TcmN, and TcmN bound to NAR-R and NAR-S.

Overall inspection of the MD trajectories revealed that NAR-R exited the TcmN catalytic cavity more frequently than NAR-S (Figure S27), whereas INT12 remained stably bound in all replicates, with few frames observed outside the cavity (Figure S29). To quantify ligand retention, we calculated the centroid of the TcmN cavity based on the centers of mass of the heavy atoms of cavity-defining residues (Figure S28 and S30), identified using the MOLE web server^38^ (mole.upol.cz). The distance between the ligand’s center of mass and the cavity centroid was then measured. By visual inspection, a threshold of 20 Å was defined as the distance beyond which the ligand was considered to have left the binding cavity. According to this criterion, ligand dissociation was observed in 19% of frames for the NAR-R 71w pose, 14% for the NAR-R 586w pose, and only 4% for the NAR-S 461w pose (Figure S9. These results indicate that the interaction of TcmN with the R-isomer of naringenin is weaker than with the S-isomer. Therefore, only the simulations performed with the NAR-S 461w pose were selected for subsequent analyses.

To investigate the conformational dynamics of free and ligand-bound TcmN, histograms of all MD simulation trajectories were analyzed. Root-mean-square deviation (RMSD) analysis showed that INT12-bound TcmN exhibited higher deviations from the initial structure compared with free and naringenin-bound TcmN, indicating greater structural rearrangements upon INT12 binding (Figure S10, S13, S16, S19, S22). The radius of gyration further revealed that INT12-bound TcmN adopted a larger overall volume than either free or naringenin-bound TcmN (Figure S12, S15, S18, S21, S24).

Cα root-mean-square fluctuation (RMSF) profiles were calculated to assess residue-level dynamics (Figure S11, S14, S17, S20, S23). All systems displayed elevated dynamics in loops L5 and L9, while INT12-bound TcmN also exhibited increased mobility in loop L3 (Figure 4A). To further investigate cavity dynamics, we measured the distance between loops L5 and L9 as a proxy for the opening and closing of the catalytic pocket. The resulting distance distributions revealed two distinct conformational populations, corresponding to closed and open states (Figure 4B). The free TcmN samples both conformations, with the open state being less populated. Upon Naringenin binding, the equilibrium shifted predominantly toward the closed state. In contrast, INT12 binding markedly favored the open conformation, indicating that, unlike the smaller ligand Naringenin, INT12 induces substantial conformational rearrangements in TcmN upon binding.

**Figure 4.**
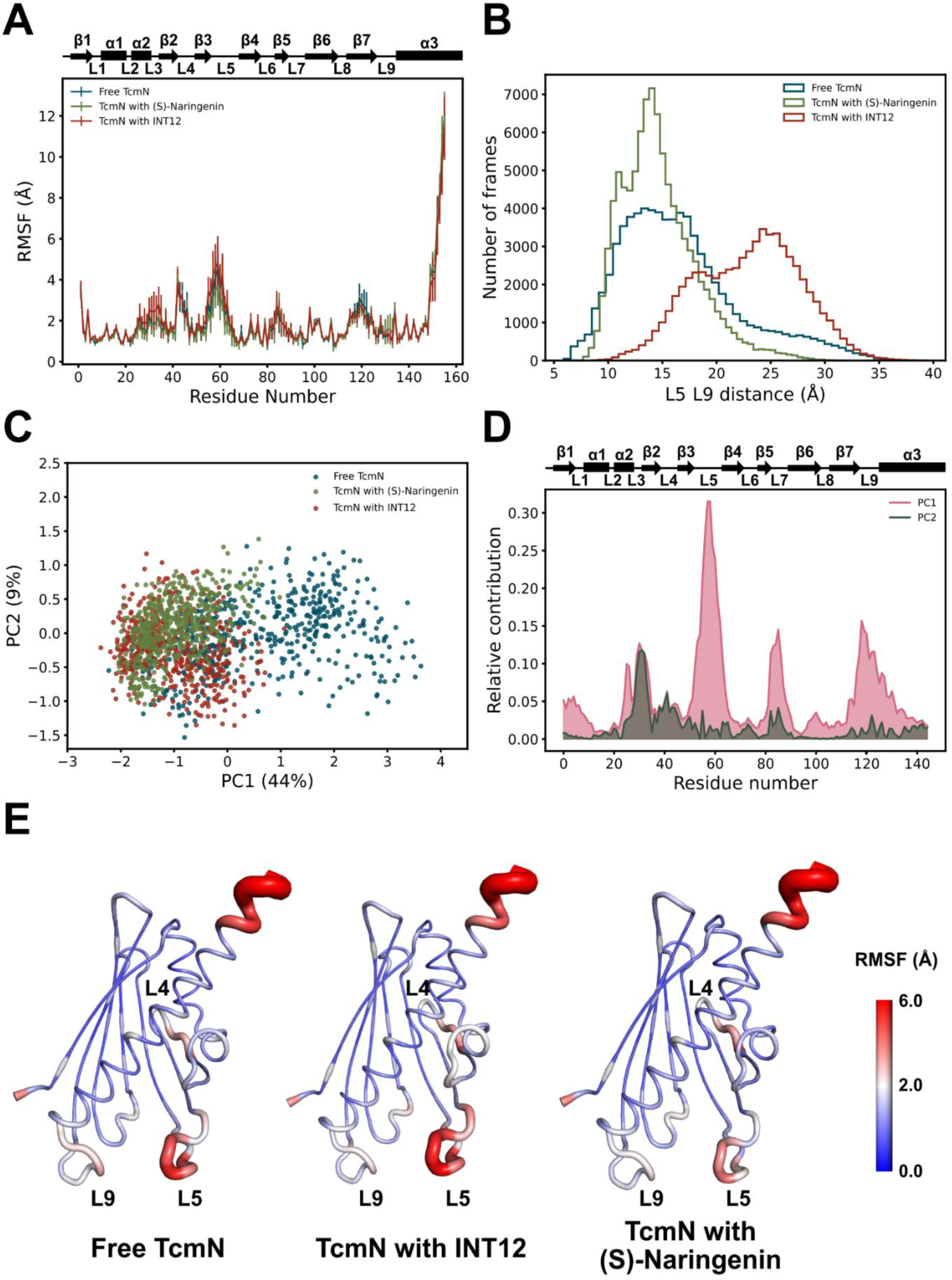
TcmN conformational diversity by MD simulations: A) Average RMSF calculated from Cα coordinates in MD simulations as a function of TcmN residue number. B) Histogram representing the distribution of distances between R61(L5) - P119(L9) from MD simulations. Combined PCA of Free TcmN, TcmN with Naringenin, and TcmN with INT12. D) Per residue contributions to PC1 and PC2 in Å. H) L5-L9 distance along PC1.E) RMSF mapped on TcmN structure.

Principal component analysis (PCA) was applied to identify dominant motions across the MD trajectories. The first two components, PC1 and PC2, accounted for 43% and 9% of the Cα variance, respectively (Figure 4C and S25). Residues from loops L3, L5, L7, and L9 at the entrance of the central cavity were the main contributors to these modes (Figure 4D). Notably, PC1 showed a strong correlation with the L5–L9 distance (Figure S26), highlighting its role in regulating the opening of the cavity (Fig. 4E).

These results demonstrate that TcmN exhibits pronounced conformational plasticity, which is essential for its catalytic function. Free TcmN preferentially adopts a closed conformation and probes the open conformation, an event probably related to substrate binding, whereas binding to linear intermediates such as INT12 promotes an open state required to accommodate the extended polyketide chain. Such conformational transitions likely facilitate sequential steps of polyketide biosynthesis by enabling substrate entry, orienting the chain for regiospecific cyclization, and allowing product release after catalysis.

Histograms of the L5–L9 loop distance show that free TcmN samples two conformational populations, corresponding to closed (μ1 ≈ 14.4 Å) and open (μ2 ≈ 24.0 Å) states (Figure 5A). In contrast, Naringenin-bound TcmN predominantly adopts a closed conformation (μ ≈ 13.9 Å), consistent with the smaller size of the ligand and the lack of need for major conformational rearrangements (Fig. 5C). Binding of INT12 shifted the equilibrium toward more open conformations, populating both intermediate (μ1 ≈ 17.4 Å) and fully open (μ2 ≈ 24.9 Å) states (Fig. 5E). Representative structures illustrate the conformational diversity of TcmN in the apo (blue), naringenin-bound (green), and INT12-bound (orange) states (Fig. 5B, 5D, 5F). These data show that while free TcmN samples both open and closed conformations, ligand binding alters this equilibrium in a ligand-dependent manner: naringenin stabilizes the closed state, whereas INT12 promotes opening of the catalytic cavity, in agreement with its requirement to accommodate a linear and polyketide chain.

**Figure 5:**
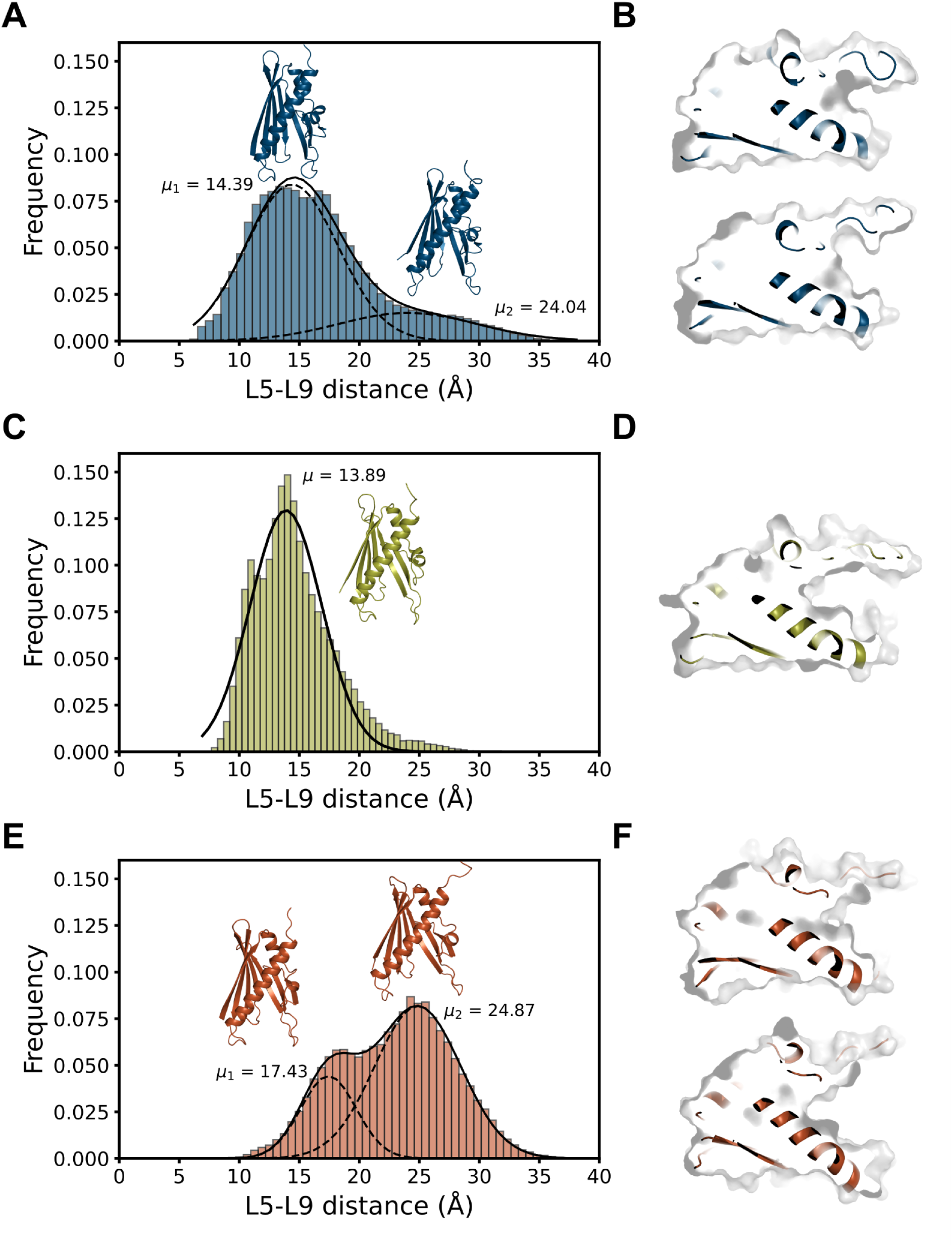
L5-L9 distance variance on MD simulations. Histogram represents the distribution of distances between the Cα of R61(L5) and P119(L9) from MD simulations of Free TcmN (A), TcmN with Naringenin (C), and TcmN with INT12 (E). B, D, and F - Ribbon representation of TcmN poses of the average distances between L5 and L9 presented in the histograms in A, C, and E, respectively.

In addition to the changes in the TcmN conformation upon binding of Naringenin and INT12, as shown in Figures 4 and 5, we also investigate the conformations of Naringenin and INT12 within the TcmN cavity. Analysis of the distance between the ligand center of mass and the cavity centroid reveals a fundamental difference in binding behavior between Naringenin and INT12, as shown in the Ligand-Cavity distance as a function of MD simulation time (Figure 6A and S27-S30). INT12 remains consistently close to the interior of the cavity, exhibiting only minor fluctuations. This stable binding is quantified by kernel density estimation (KDE) of the Ligand–Cavity distance in Figure 6B and 6C, which shows a major population at a close distance (3 Å) from the cavity centroid, with a minor population at a more distant position (7 Å). The structural basis for this stability is visualized in Figure 6D, which shows INT12 deeply buried in the cavity (INT12 conformation colored in red). The minor IN12 sampled conformation is shown in Figure 6D, in blue.

**Figure 6.**
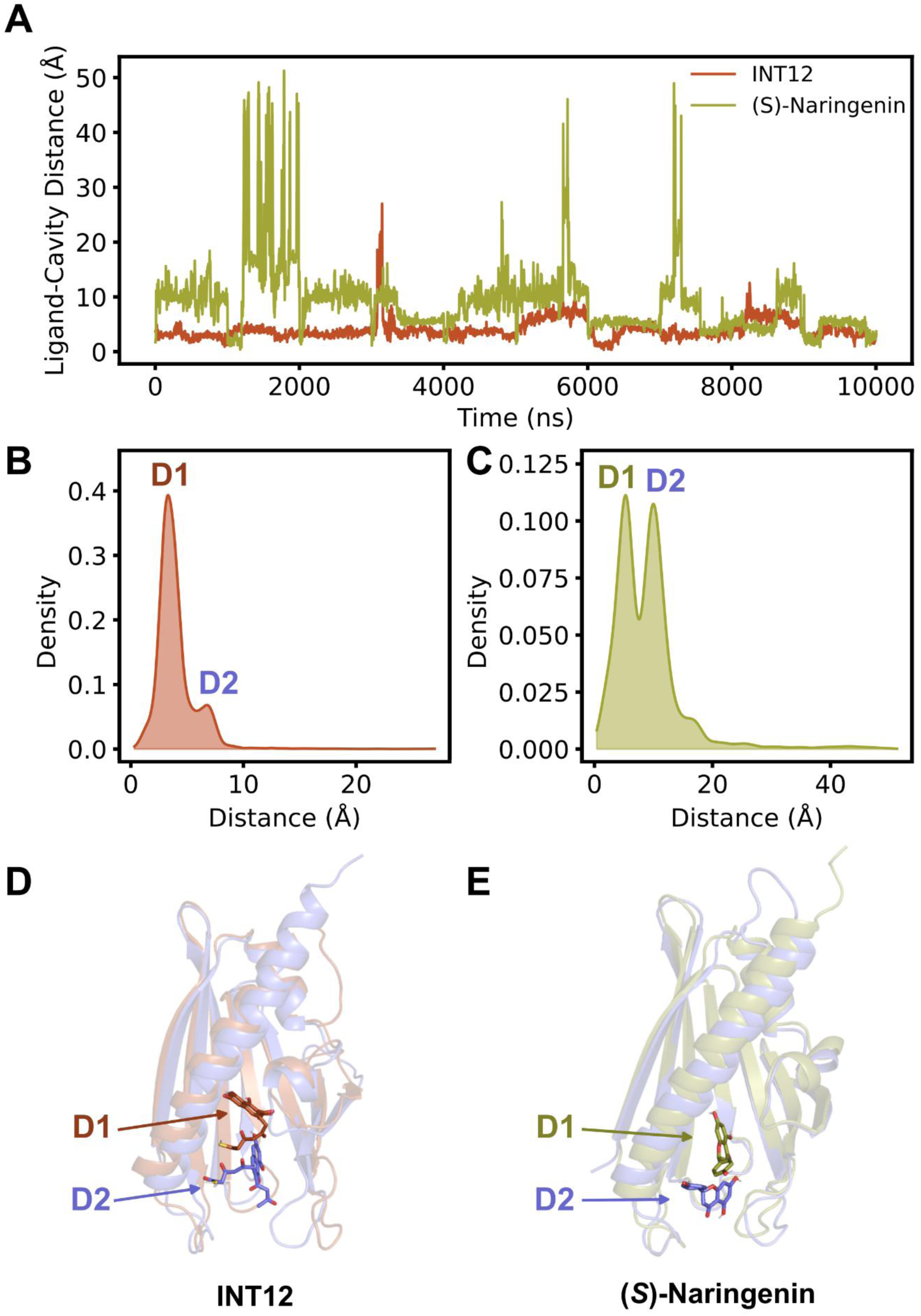
Ligand–cavity distance in MD simulations. A) Time evolution of the distance between the center of mass of INT12 and (S)-Naringenin and the centroid of the TcmN cavity. B) Kernel density estimation (KDE) of the distance distribution for INT12. C) KDE of the distance distribution for (S)-Naringenin. D) Ribbon representation of two representative conformations of INT12 within the TcmN cavity (the deep conformation shown in orange and the distal in blue). E) Ribbon representation of two representative conformations of (S)-Naringenin within the TcmN cavity (the deep conformation shown in green and the distal in blue).

(S)-naringenin samples two distinct conformational states, as indicated by its fluctuating distance profile (Figure 6A) and by the bimodal KDE plot (Figure 6C). One population corresponds to a pose buried in the cavity interior, with a Ligand-Cavity distance of ∼5 Å. In contrast, a second, similarly probable population is located at a greater distance from the centroid (10 Å). This second conformation of Naringenin, farther from the cavity center, is illustrated in Figure 6E, and is closer to the flexible loops that form the entrance to the binding site.

Figure 7 shows representative structures of INT12 and (S)-Naringenin bound to the enzyme in both proximal and distal conformations. In the first conformation, INT12 is positioned deep within the catalytic cavity, engaging residues such as W65, F88, and T132 in the narrow neck region. This supports the proposal by Ames et al.^3^ that this constricted area limits the positional and conformational freedom of the bound polyketide intermediate. Their mutational analysis further identified R69 as essential for catalyzing the C9-C14 ring pattern. Additionally, W108, located deep within the cavity, appears critical for proper folding, cyclization, and aromatization of the intermediates, likely by contributing to the overall structural stability of these processes.

**Figure 7.**
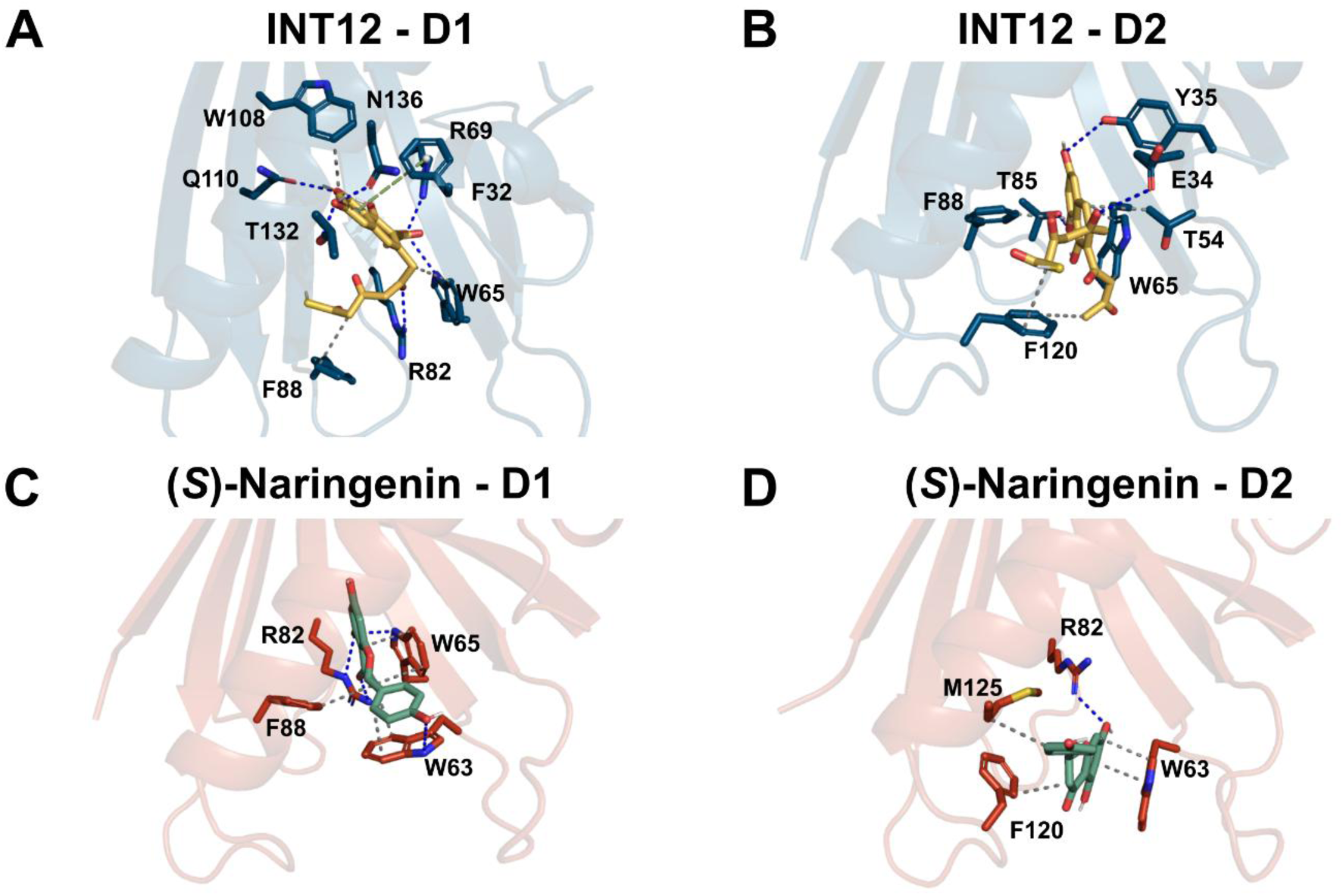
Protein-ligand interactions in proximal and distal conformations. (A) INT12 interactions in the proximal D1 conformation. (B) INT12 interactions in the distal D2 conformation. (C) (S)-Naringenin interactions in the proximal D1 conformation. (D) (S)-Naringenin interactions in the distal D2 conformation.

In contrast, the first conformation of (S)-Naringenin is positioned closer to the cavity entrance, where it interacts with W63, W65, R82, and F88. A second, distal conformation was also observed, situated even nearer to the entrance loops and interacting with W63, R82, F120, and M125. These structural findings are consistent with CSP analysis. Significant CSPs were detected in residues surrounding the TcmN cavity and alpha-helix 3, including T5, S33, Y35, D57, W63, F120, T126, N136, and M137. Notably, residues F120 and W63—located in loops near the cavity entrance—exhibited some of the most pronounced shifts, corroborating their role in ligand binding.

Hydrogen bond occupancy analysis performed with VMD (occupancy >5% for INT12; >3% for (S)-Naringenin) revealed distinct interaction networks for the two ligands (Figure 8). For INT12, the key interacting residues are R82, T85, Q110, and T133. These residues are located near loop 7 and within a narrow constriction of the TcmN cavity, effectively locking the ligand in a productive pose. For (S)-Naringenin, the major hydrogen-bonding residues identified were E34, R82, Q110, and T133.

**Figure 8.**
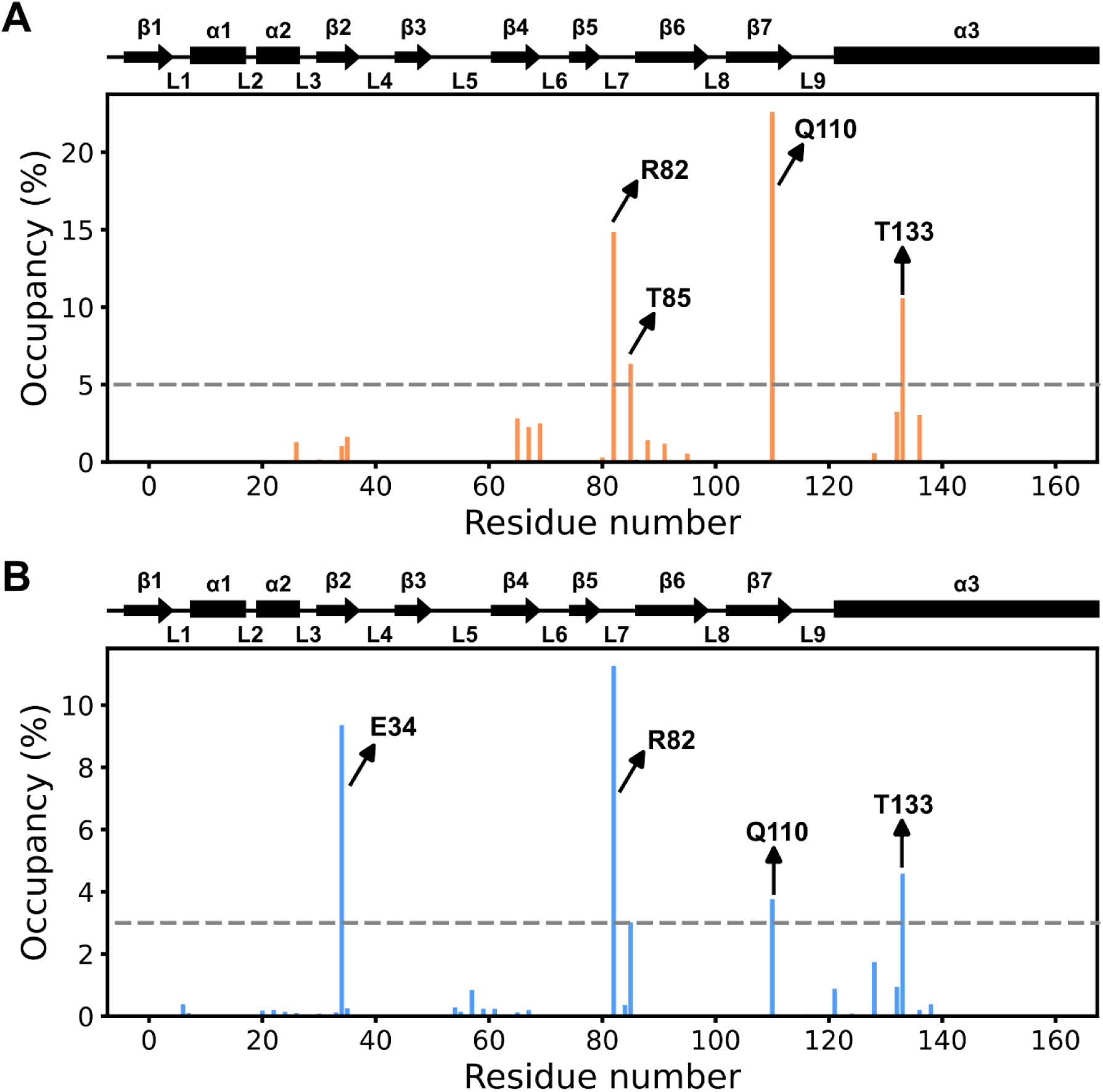
Occupancy of hydrogen bonds between TcmN residues and the ligand. (A) INT12. (B) (S)-Naringenin.

Our findings provide direct structural support for previously proposed catalytic mechanisms. Ames et al.^3^ identified R82 as highly conserved in ARO/CYC enzymes and proposed its critical role in second-ring cyclization and aromatization. Furthermore, mutational studies on the TcmN homolog WhiE, conducted by Lee et al.^4^, demonstrated that R82 is essential for enzyme activity and that E34 may coordinate an active water molecule during catalysis. Notably, Q110, a key residue defining the cavity bottleneck, has been implicated in C9-C14 cyclization, potentially facilitating this step through hydrogen bonding to intermediates or by positioning catalytic water molecules. The persistent hydrogen bonds observed between these catalytically crucial residues (R82, E34, Q110) and the intermediate INT12 and its analog (S)-Naringenin corroborate their functional importance.

For residues that formed frequent hydrogen bonds, we identified the specific ligand atoms involved in these contacts (distance < 3.5 Å, Figure S31 The interactions were highly specific and provided a mechanistic explanation for the differential stability of the two complexes.

For INT12, the interactions were precise and localized. The critical residue R82 interacted predominantly with the oxygen atoms of the ketone carbonyls on the polyketide tail and may adjust the orientation of the ligand. Simultaneously, Q110, T85, and T133 formed hydrogen bonds primarily with atoms on the first aromatic ring. This interaction pattern suggests that the two meta-positioned hydroxyl groups on this ring are crucial for molecular recognition and for positioning the ligand in a productive orientation within the catalytic site.

For (S)-Naringenin, the binding mode was less specific. While R82 also contacted the ligand, its interactions were distributed across the flavonoid ring hydroxyl and ketone groups. Similarly, E34, Q110, and T133 interacted with a broader set of atoms, suggesting a more dynamic, less rigid binding pose. This latter observation provides a structural rationale for our experimental NMR data, in which the NMR signals for these same residues (E34, Q110, T133) disappeared upon titration with (S)-naringenin. The loss of signal is highly suggestive of a significant conformational change or an intermediate-timescale exchange process induced by ligand binding, consistent with the dynamic, less-specific interaction network observed in the MD simulations.

As shown in Figures 7C and 7D, R82 adopts markedly distinct conformations in the two binding modes of (S)-naringenin, suggesting that this residue may play multiple functional roles during catalysis (Figure S32). This conformational plasticity is consistent with our NMR data: the ¹⁵N–HSQC spectrum of TcmN shows no detectable signal for R82, indicating that, even in the ligand-free state, this residue undergoes conformational exchange on an intermediate chemical-shift timescale. Such dynamic behavior supports the notion that R82 acts as a flexible catalytic switch, sampling distinct rotameric states required for different stages of the reaction.

Overall, these results demonstrate that TcmN exhibits pronounced conformational plasticity, which is essential for its catalytic function. Free TcmN preferentially adopts a closed conformation, whereas binding to linear intermediates such as INT12 promotes an open state required to accommodate the extended polyketide chain. Such conformational transitions likely facilitate sequential steps of polyketide biosynthesis by enabling substrate entry, orienting the chain for regiospecific cyclization, and allowing product release after catalysis.

## CONCLUSIONS

In this work, we provide structural and dynamic evidence that conformational plasticity is essential for TcmN function. By integrating NMR spectroscopy, calorimetry, docking, and long-timescale molecular dynamics simulations, we demonstrate that apo TcmN undergoes dynamic exchange between open and closed conformations, and that ligand identity determines the equilibrium distribution. Naringenin stabilizes a compact, closed state, consistent with substrate protection from solvent exposure. In contrast, the tetracenomycin C intermediate INT12 drives the enzyme toward expanded, open states that facilitate the accommodation of linear intermediates and subsequent regiospecific cyclization. These findings support a ligand-gated breathing model for polyketide aromatase/cyclases. Importantly, the demonstration that different intermediates bias distinct conformational ensembles highlights how sequential ligand binding choreographs catalysis and product release, while reducing the risk of aggregation of hydrophobic surfaces. Together, our results suggest that W63 functions as a gatekeeper residue controlling ligand access to the catalytic pocket, while the presence of two distinct binding sites—one core and one entrance-region—supports a sequential. This study demonstrates how combining experimental NMR with atomistic simulations can elucidate transient conformational equilibria and reveal the molecular strategies underlying enzyme adaptability. Beyond providing mechanistic insight into polyketide biosynthesis, our results underscore the potential of exploiting conformational dynamics to guide the rational design and engineering of new polyketide scaffolds with biomedical relevance.

## Acknowledgments

This work was supported by the Brazilian Innovation and Research Funding Agency, FINEP, grant MCTI/FINEP/FNDCT 01/2016/01.16.0050.00, by “Fundação de Amparo à Pesquisa do Estado de Minas Gerais” (FAPEMIG), grants DEMANDA UNIVERSAL/2022/APQ-00428-22 and INSTALAÇÃO MULTIUSUÁRIOS/2024/APQ-02819-24. We would like to thank CNPq (Conselho Nacional de Desenvolvimento Científico e Tecnológico) and CAPES (Coordenação de Aperfeiçoamento de Pessoal de Nível Superior) for funding support. V. S. V. was supported by CNPq grant 200611/2022-4. We also would like to thank the Laboratório de Ressonância Magnética Nuclear de Alta Resolução, LAREMAR, at UFMG, Universidade Federal de Minas Gerais, the Centro Nacional de Ressonância Magnética Nuclear at UFRJ, Universidade Federal do Rio de Janeiro, and the NMR facilities at Dana-Farber and Harvard Medical School for NMR spectrometer time, as well as the Laboratório Multiusuário do Departamento de Biofísica of the Federal University of São Paulo, for DSC spectrometer time.

## Data and Software Availability statement

All data associated with this work are publicly available in a Zenodo repository at https://doi.org/10.5281/zenodo.17896225. The repository contains molecular docking results and molecular dynamics (MD) simulation data organized into two main components: docking and MD trajectories. The docking data (docking.tar.gz) include docking poses of TcmN with intermediate 12 (INT12), (R)-naringenin (poses 71w and 586w), and (S)-naringenin, which were used to select starting structures for molecular dynamics simulations. The MD data are provided in TcmN-Trajectories.tar.gz and in separate replica archives. These include simulations of apo TcmN, TcmN bound to INT12, TcmN bound to (R)-naringenin (poses 71w and 586w), and TcmN bound to (S)-naringenin. For each system, a combined trajectory file (*-combtraj-sampled.xtc) containing a 5% regularly sampled trajectory, merged from ten independent 1-µs replicas, is provided, along with the corresponding topology file (*.tpr). In addition, individual production trajectories for each replica are available as separate compressed archives (*-reps.tar.gz). All simulations were performed using GROMACS.

## AUTHOR INFORMATION

### Corresponding Author

**Adolfo H. Moraes** – Departamento de Química, Instituto de Ciências Exatas, Universidade Federal de Minas Gerais, Av. Antonio Carlos 6627, 31270-901 Belo Horizonte, MG, Brazil.

### Author Contributions

The manuscript was written through the contributions of all authors. All authors have given approval to the last version of the manuscript.

