## Supporting Information for "The Role of Conformational Changes in TcmN Aromatase/Cyclase in Polyketide Biosynthesis"

**Table S1.** Final concentrations of TcmN and naringenin at each titration point in the $^{1}H$–$^{15}N$ HSQC experiments acquired at 600 MHz. Protein dilution due to the addition of ligand solution in DMSO-$d_6$ was considered.

| TcmN concentration (μM) | Naringenin concentration (μM) | [L]:[P] ratio (Naringenin/TcmN) |
|---|---|---|
| 80.0 | 0 | 0 |
| 79.87 | 41.6 | 0.52 |
| 79.74 | 83.2 | 1.04 |
| 79.61 | 124.8 | 1.57 |
| 79.35 | 208 | 2.62 |
| 78.72 | 416 | 5.28 |
| 78.28 | 561.6 | 7.2 |

**Table S2.** Final concentrations of TcmN and naringenin at each titration point in the $^{1}H$–$^{15}N$ HSQC experiments acquired at 800 MHz. Protein dilution due to the addition of ligand solution in DMSO-$d_6$ was considered.

| TcmN concentration (μM) | Naringenin concentration (μM) | [L]:[P] ratio (Naringenin/TcmN) |
|---|---|---|
| 50.00 | 0 | 0 |
| 49.98 | 25 | 0.50 |
| 49.95 | 50 | 1.00 |
| 49.9 | 100 | 2.00 |
| 49.8 | 200 | 4.02 |
| 49.6 | 400 | 8.06 |
| 49.31 | 1000 | 20.28 |
| 48.83 | 2000 | 40.96 |

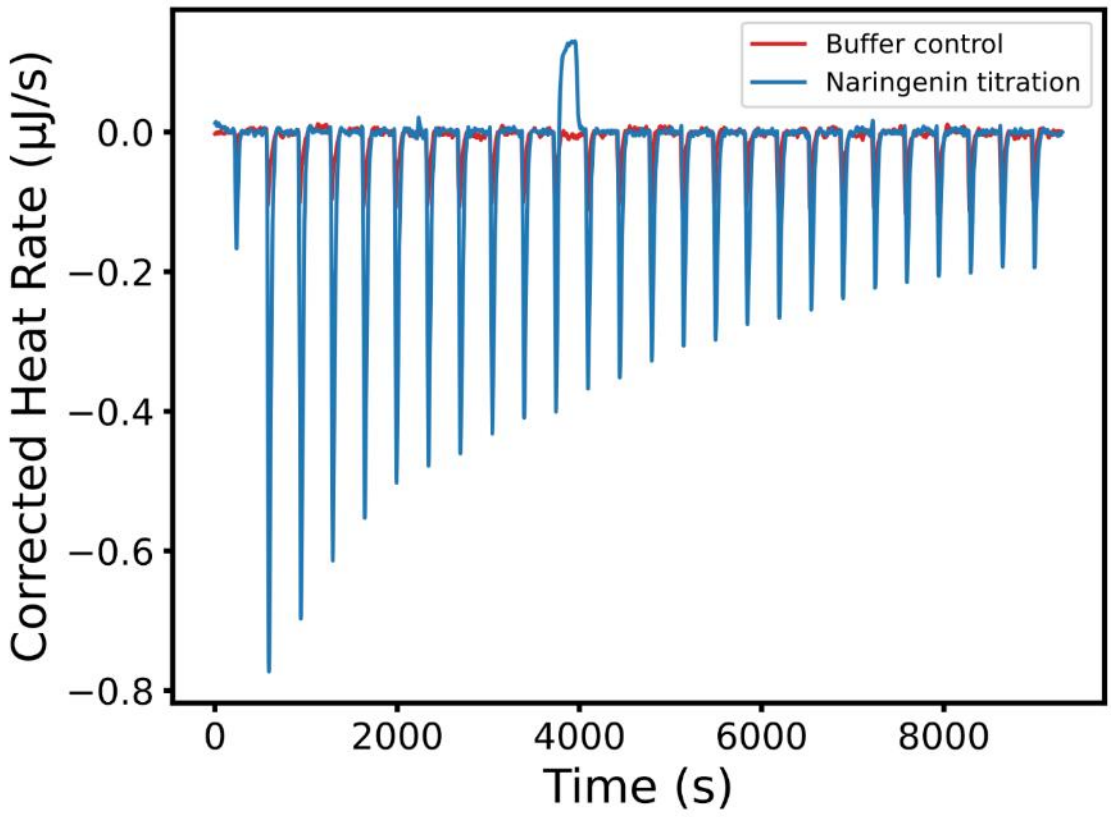


**Figure S 1: ITC experiment TcmN's reaction intermediate and product analog naringenin.** Calorimetric titration of Naringenin into a sample cell containing TcmN. The titration was performed at 25 °C in 20 mM sodium phosphate pH 7, 100 mM NaCl, and 4% DMSO.

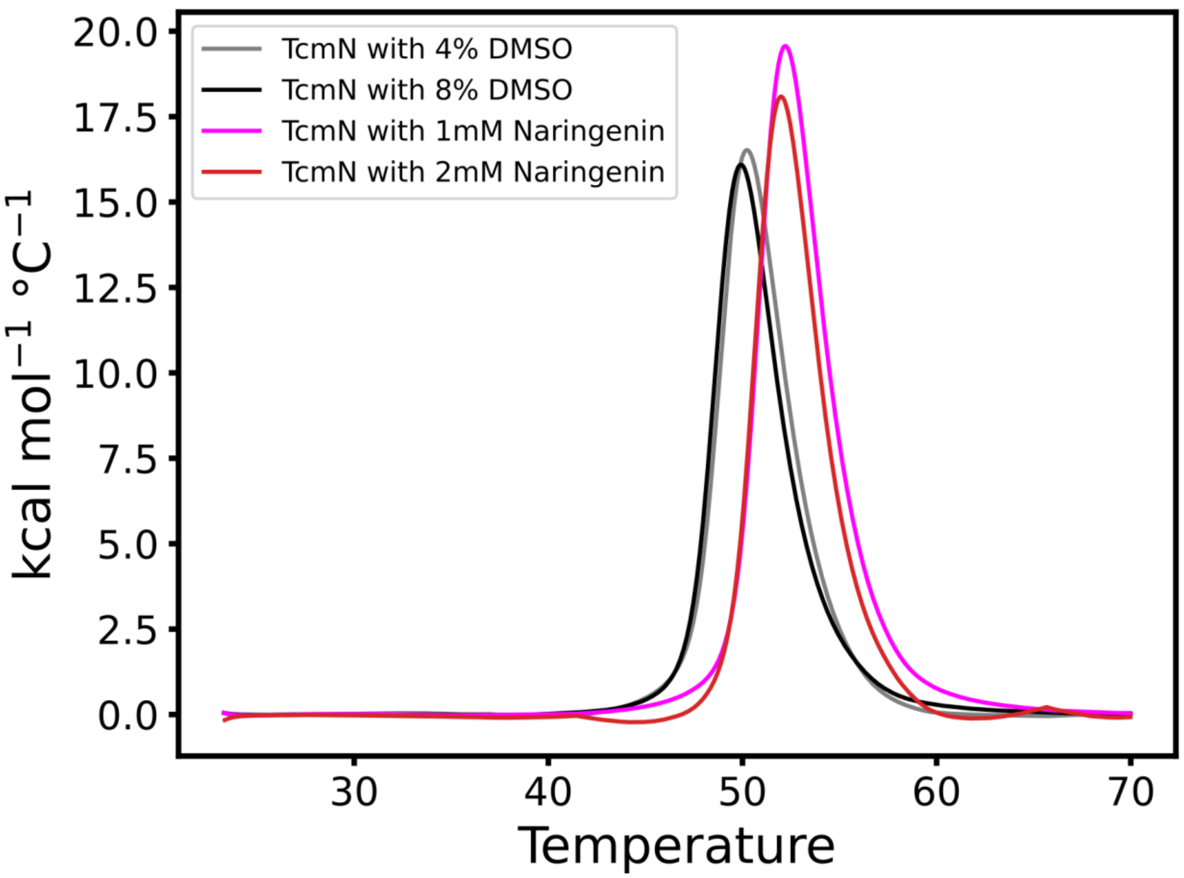


**Figure S 2 - Effect of Naringenin and DMSO on TcmN thermostability.** Overlayed experimental thermograms of 211 µM TcmN in 20 mM sodium phosphate pH 7, 100 mM NaCl in the presence of 1 mM and 2 mM of Naringenin, and with 4 and 8% of DMSO.

.

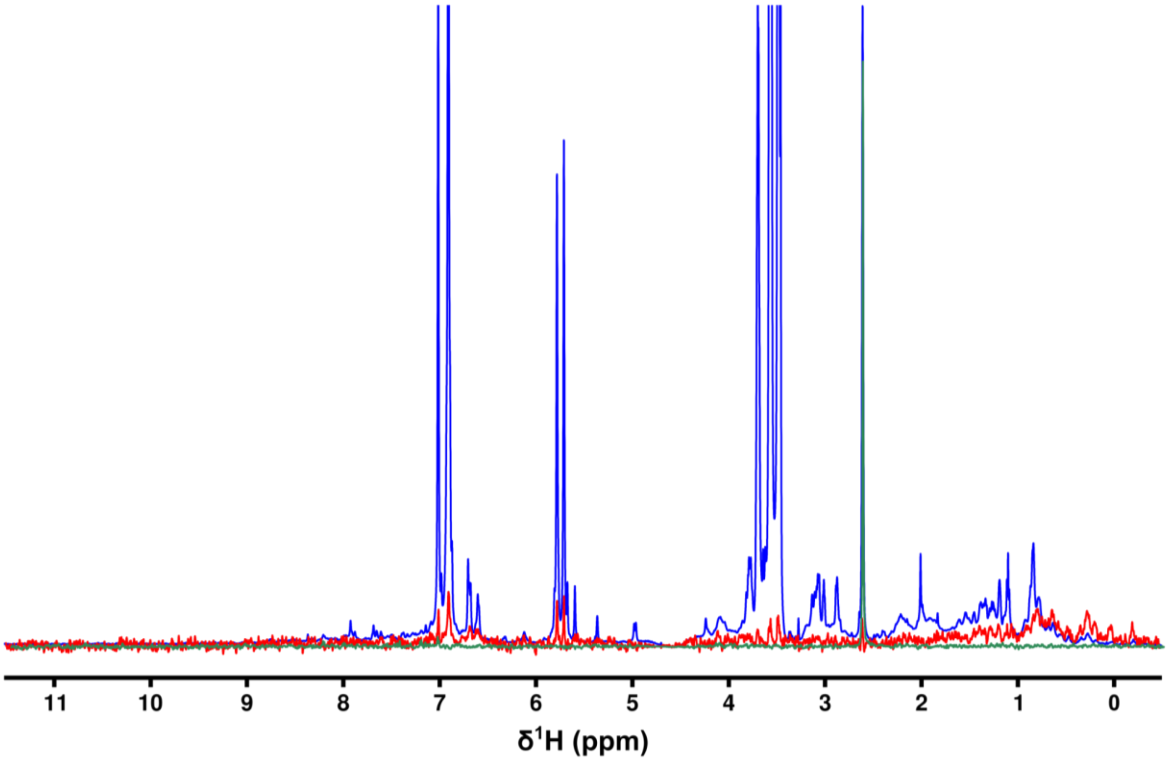


**Figure S 3 - Interaction of TcmN with taxifolin by saturation transfer difference (STD) NMR experiment.** (Blue spectrum) $^{1}$H 1D NMR spectrum of taxifolin 1 mM, TcmN 10 micromolar. (Red spectrum) STD spectrum of taxifolin 1 mM in TcmN 10 micromolar. (Green spectrum, control experiment) STD of taxifolin 1 mM. The experiments were performed at 25 °C in phosphate buffer 20 mM, NaCl 150 mM, and pH 7.0.

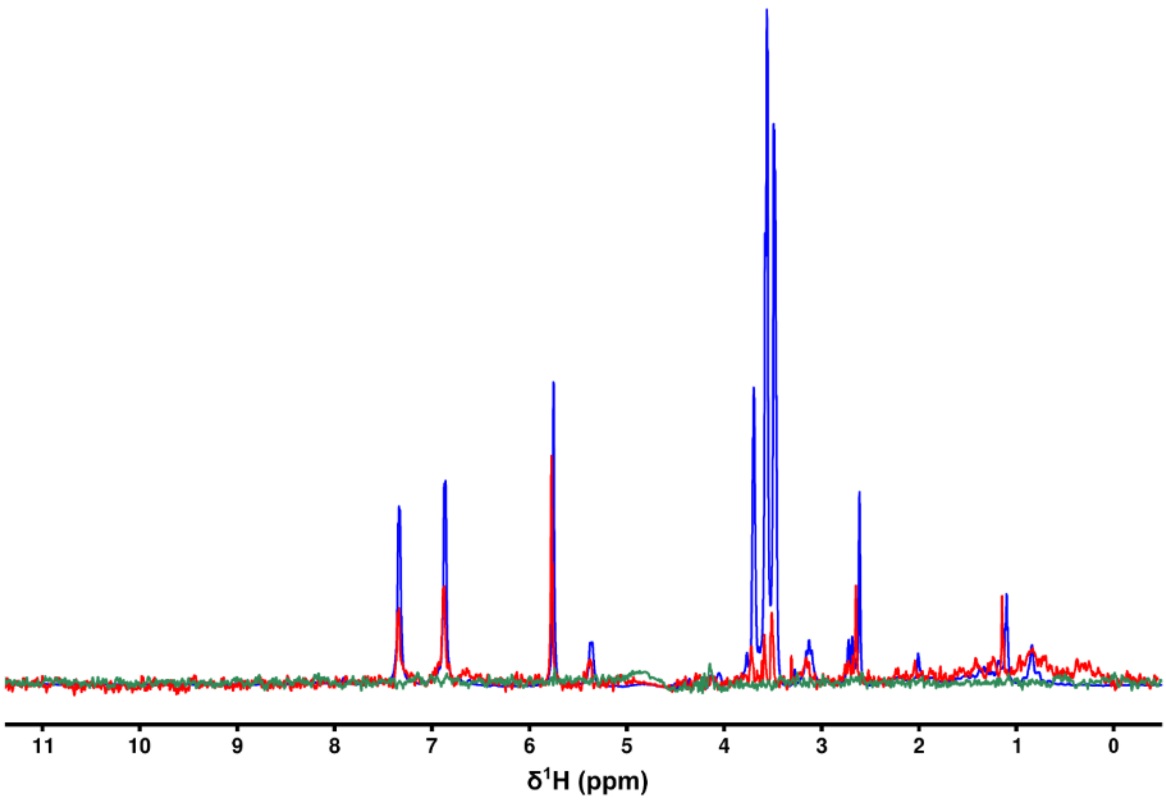


**Figure S 4 - Interaction of TcmN with naringenin by saturation transfer difference (STD) NMR experiment.** (Blue spectrum) $^{1}$H 1D NMR spectrum of naringenin 1 mM, TcmN 10 micromolar. (Red spectrum) STD spectrum of Naringenin 1 mM in TcmN 10 micromolar. (Green spectrum, control experiment) STD of Naringenin 1 mM. The experiments were performed at 25 °C in phosphate buffer 20 mM, NaCl 150 mM, and pH 7.0.

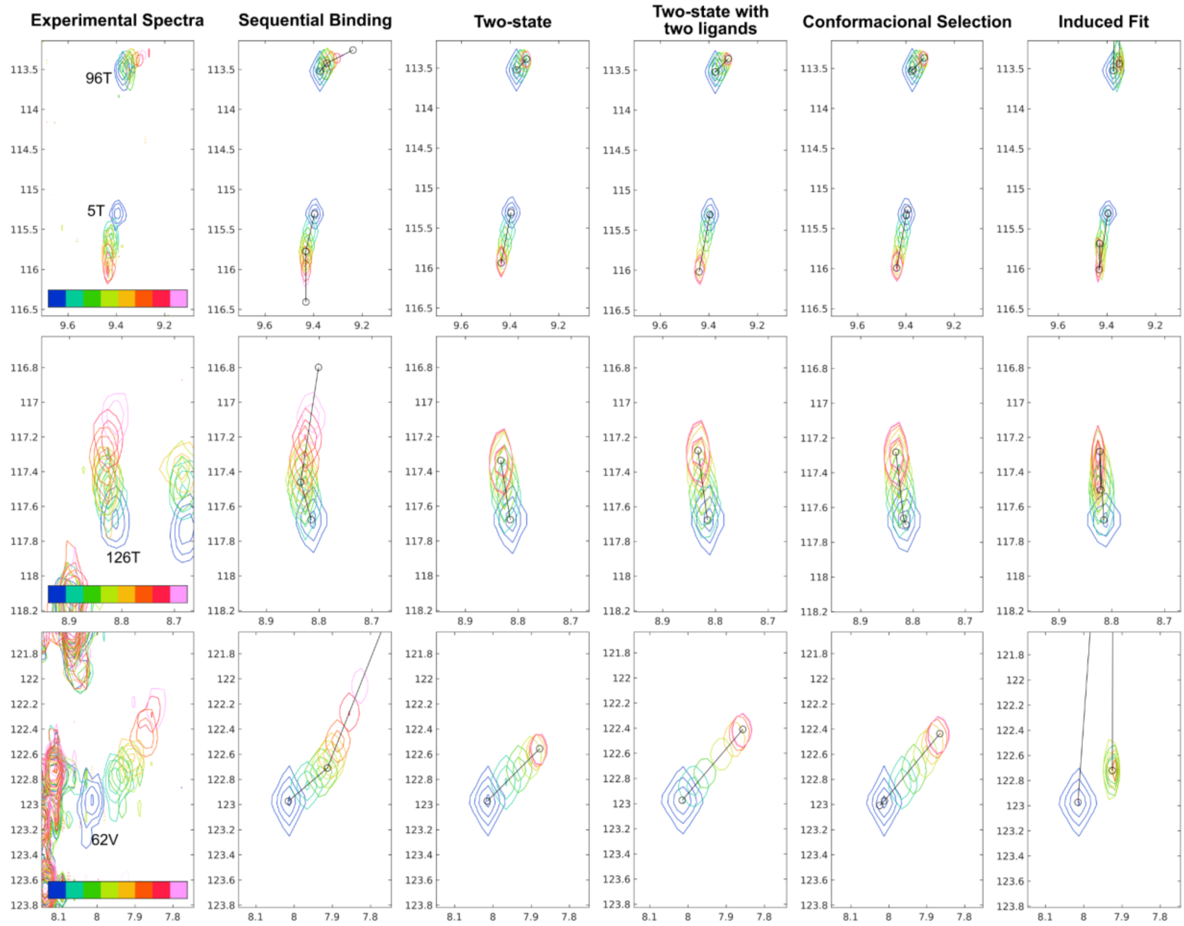


**Figure S 5 Comparison of the spectra from the $^{1}H$-$^{15}N$ HSQC experiments and the simulations performed by TITAN.** Comparison of the experiments acquired at 800 MHz and the simulations performed by TITAN for the different binding models.

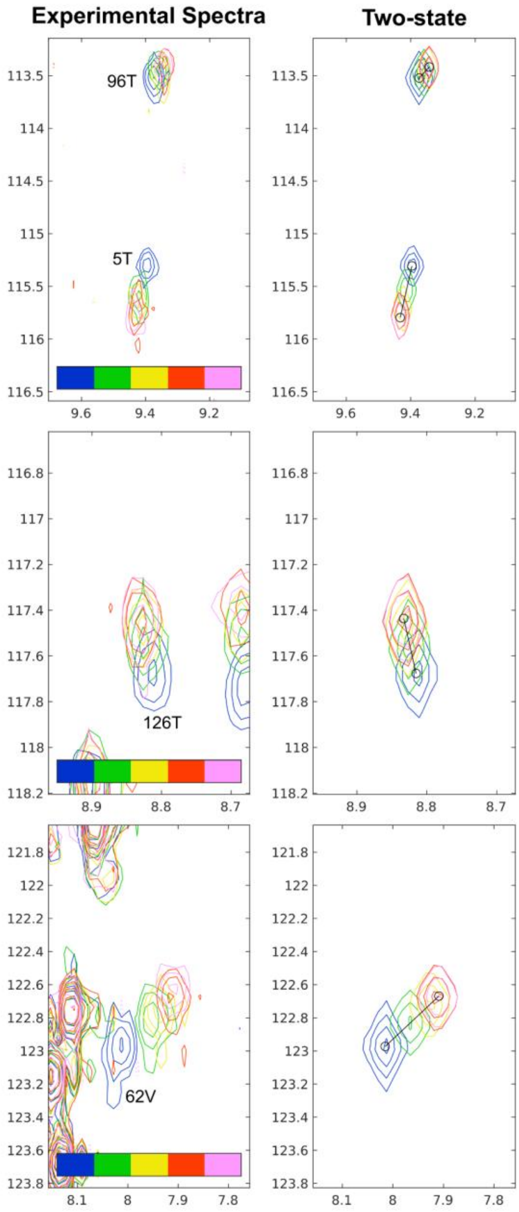


**Figure S 6 - Comparison of the spectra from the $^{1}H$-$^{15}N$ HSQC experiments and the simulations performed by TITAN.** Comparison of the experiments acquired at 800 MHz up to the titration of 200 µM of naringenin and the simulation performed by TITAN for the lock-and-key model.

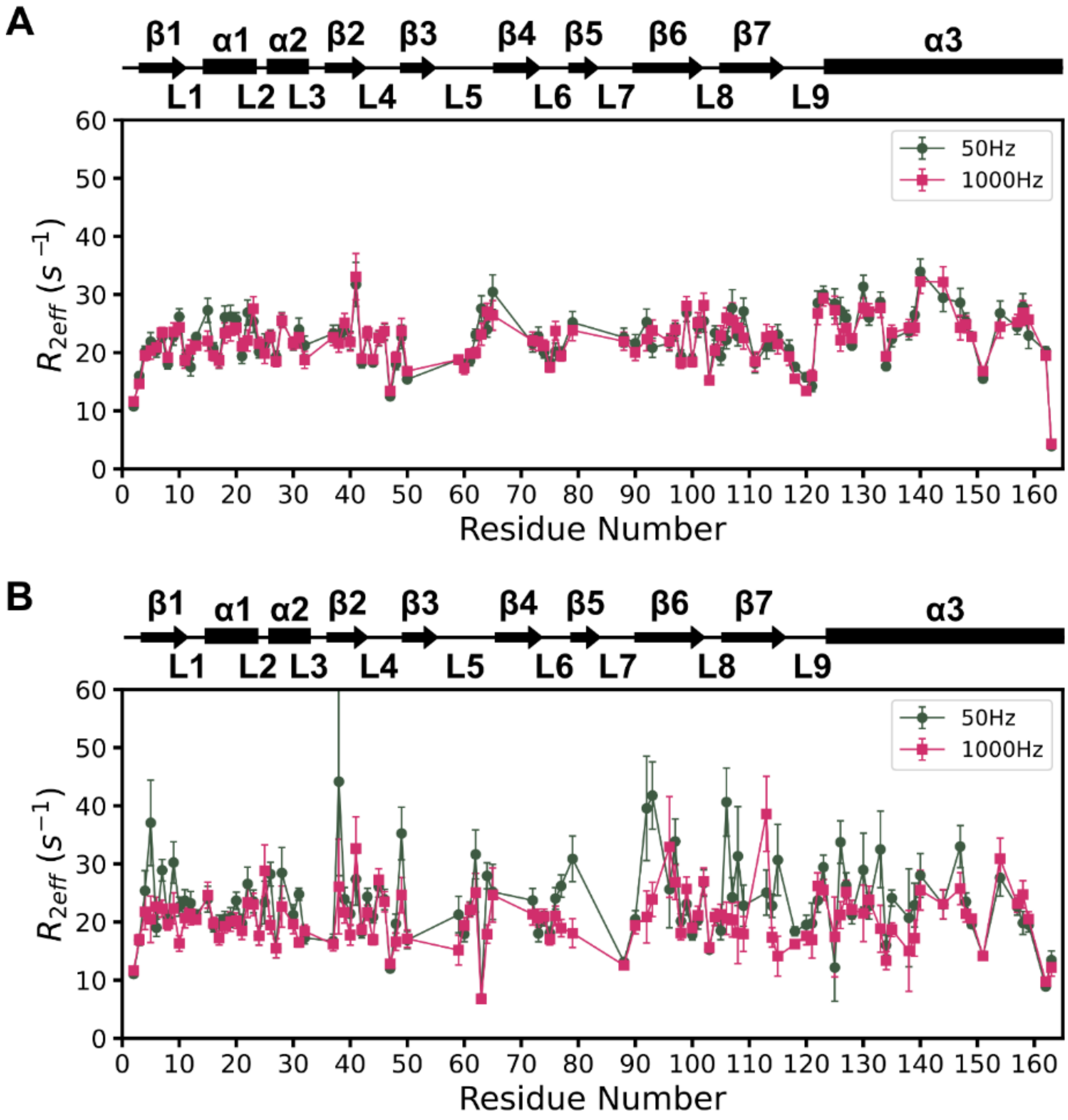


**Figure S 7 - Backbone dynamics of TcmN free and bound to naringenin by NMR spectroscopy.** (A) $R_{2,eff}$ $(s^{-1})$ rate as a function of TcmN residue number of 130 µM TcmN in 20 mM sodium phosphate, pH 7, 100 mM NaCl. (B) $R_{2,eff}$ $(s^{-1})$ rate as a function of TcmN residue number of 126 µM of TcmN titrated with 1.2 mM of Naringenin. Black squares and red circles refer to $^{15}$N-RD-CPMG relaxation dispersion experiments acquired with CPMG frequencies of 50 Hz and 1000 Hz, respectively.

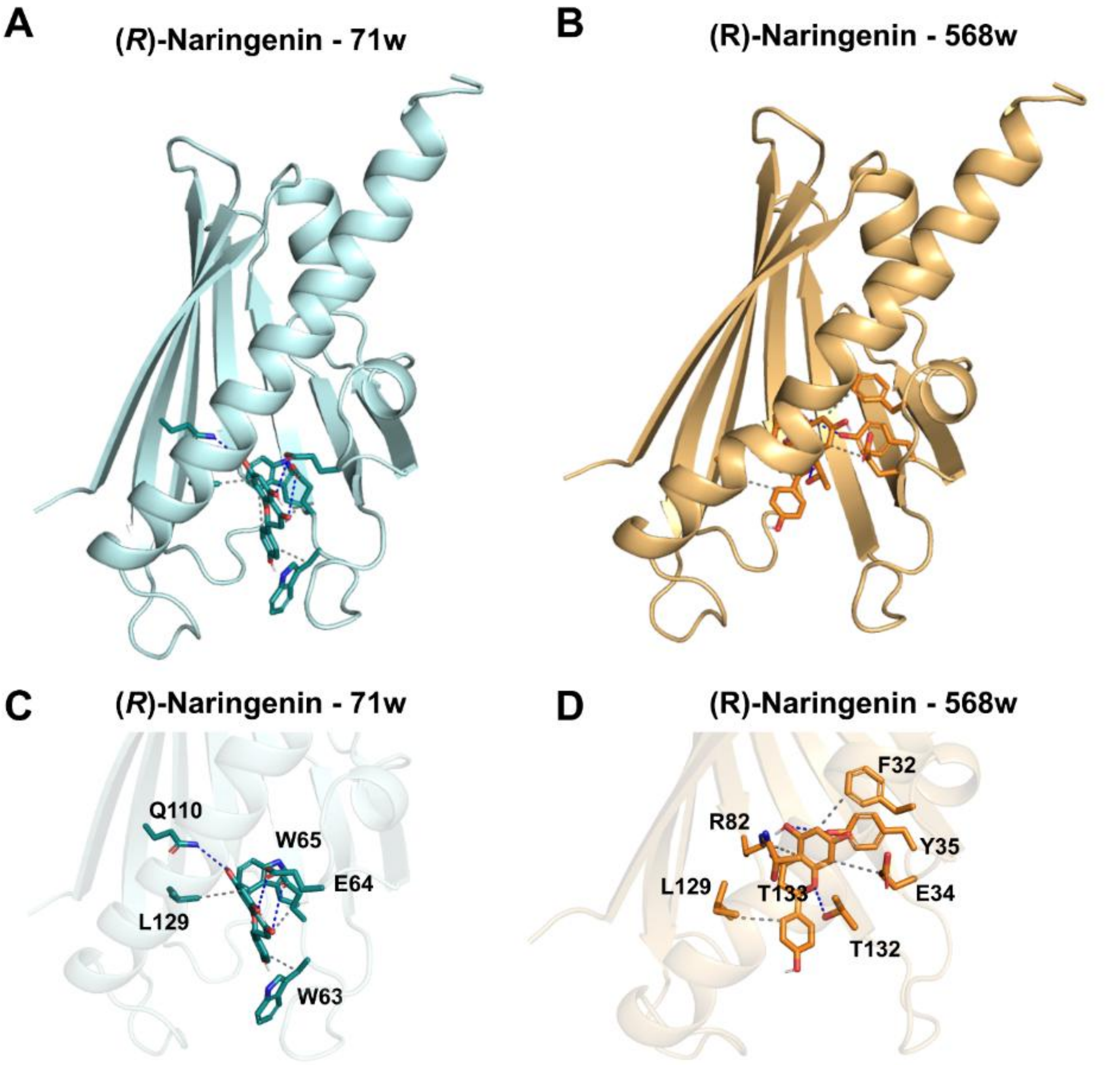


**Figure S 8 – (R)-Naringenin docking poses.** A) 71w pose. B) 568w pose. (C, D) Corresponding 2D ligand interaction diagrams detailing the specific molecular interactions (e.g., hydrogen bonds, hydrophobic contacts) for the 71w and 568w complexes, respectively.

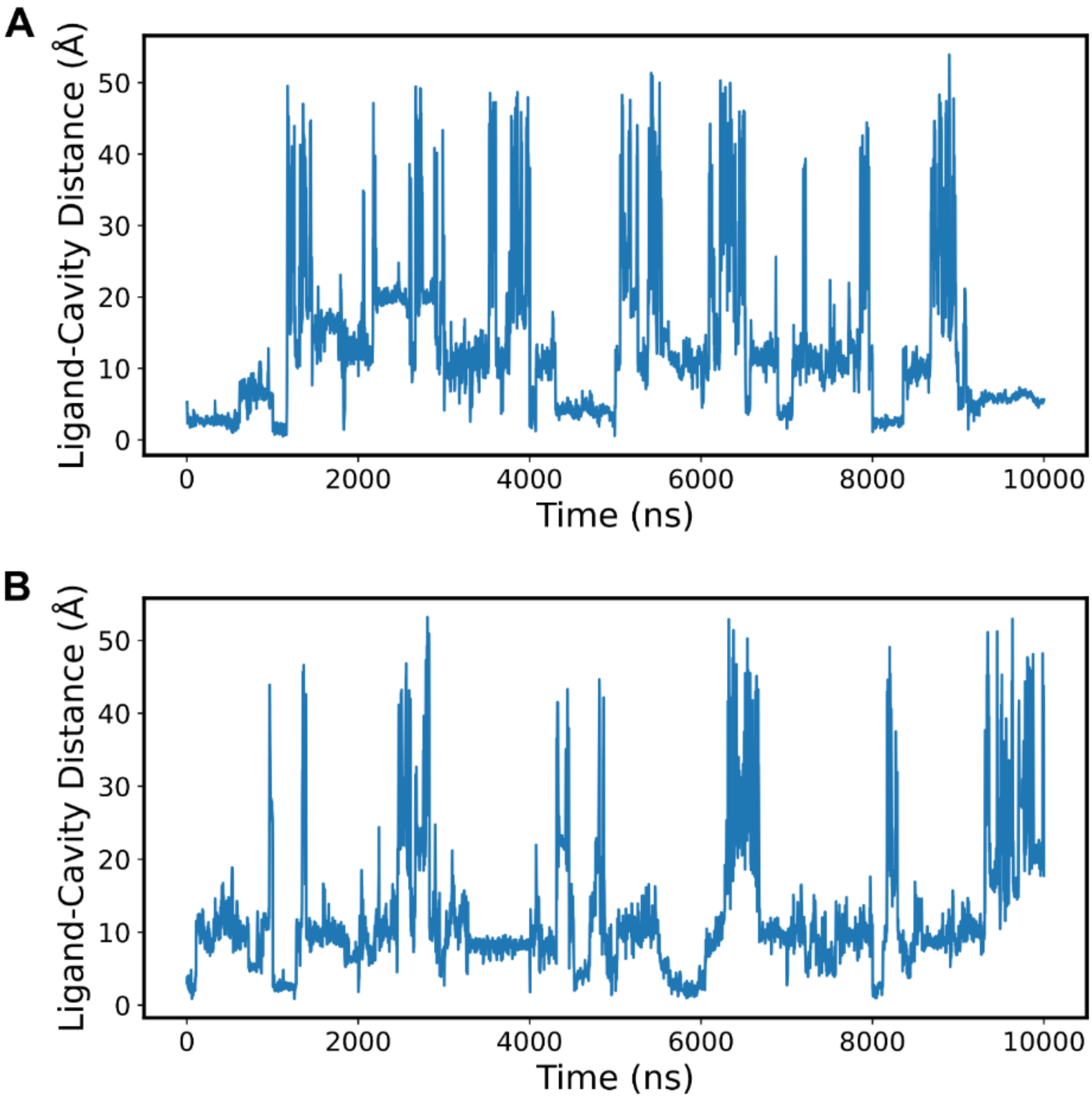


**Figure S 9 – (R)-Naringenin distance from cavity centroid.** A) MD trajectory using 71w pose. B) MD trajectory using 568w pose.

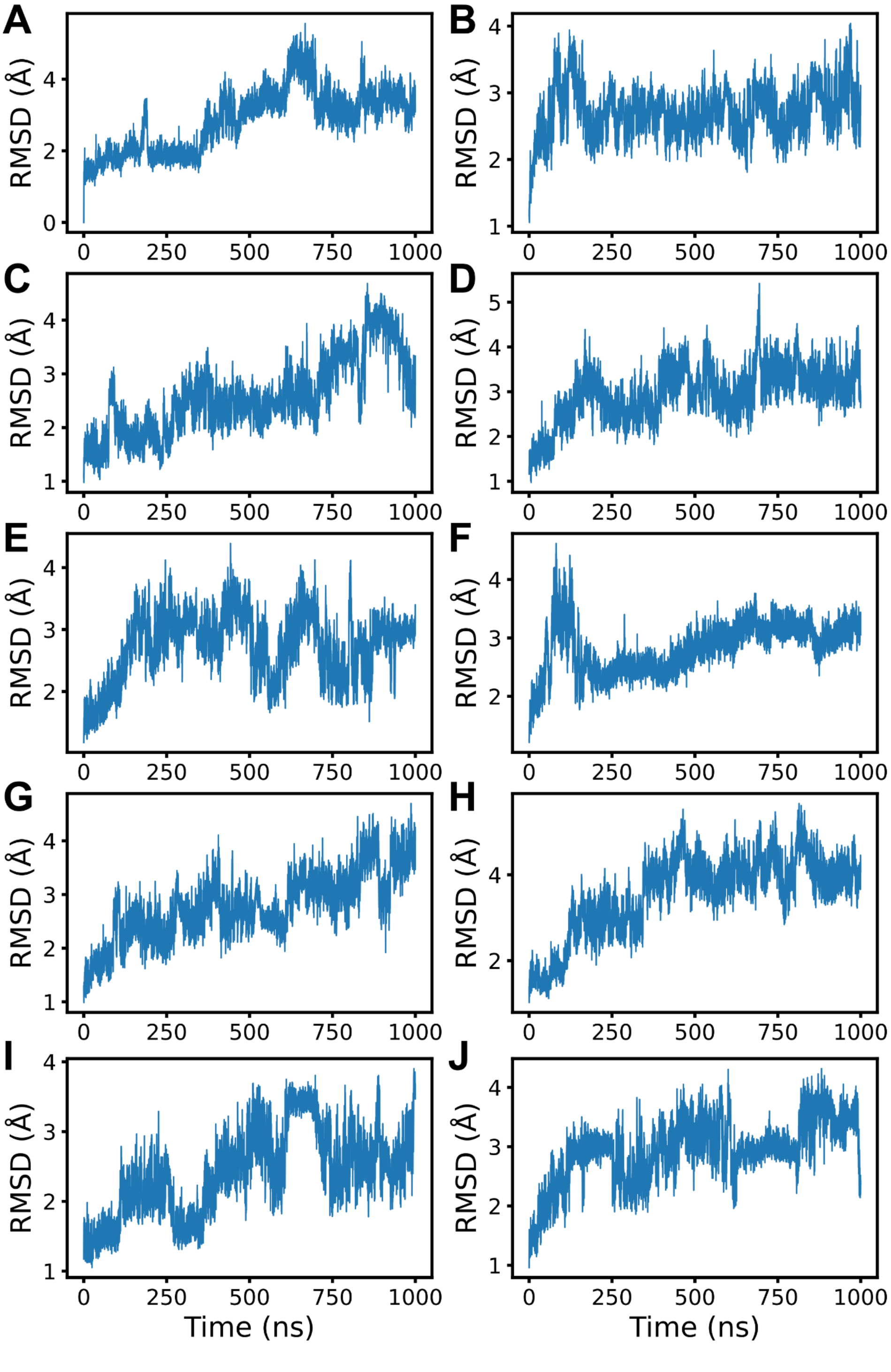


**Figure S 10 - Free TcmN molecular dynamics simulations.** Root mean square deviation (RMSD) of the MD trajectory for each replicate.

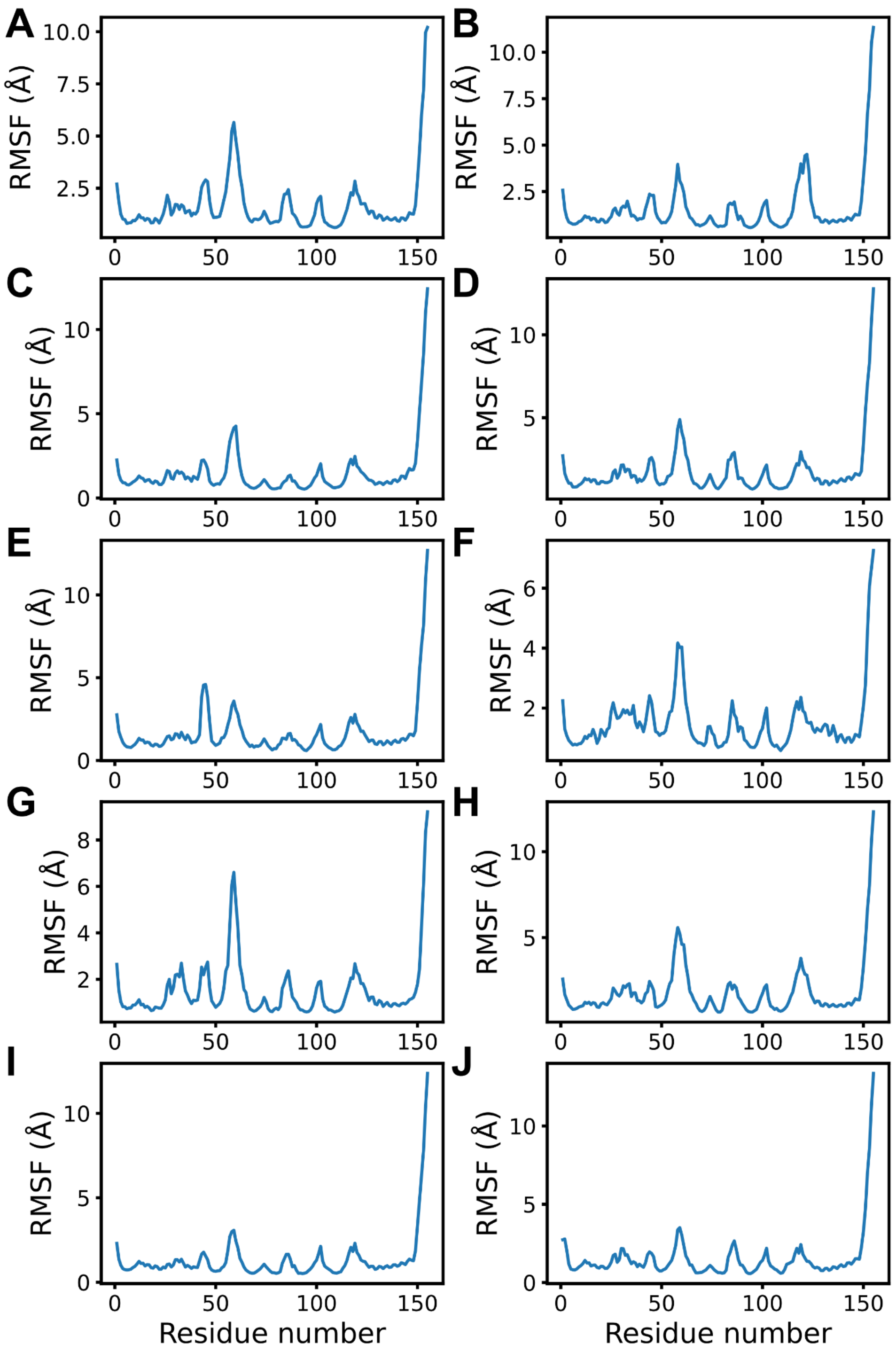


**Figure S 11 - Free TcmN molecular dynamics simulations**. Root mean square fluctuation (RMSF) of the MD trajectory for each replicate.

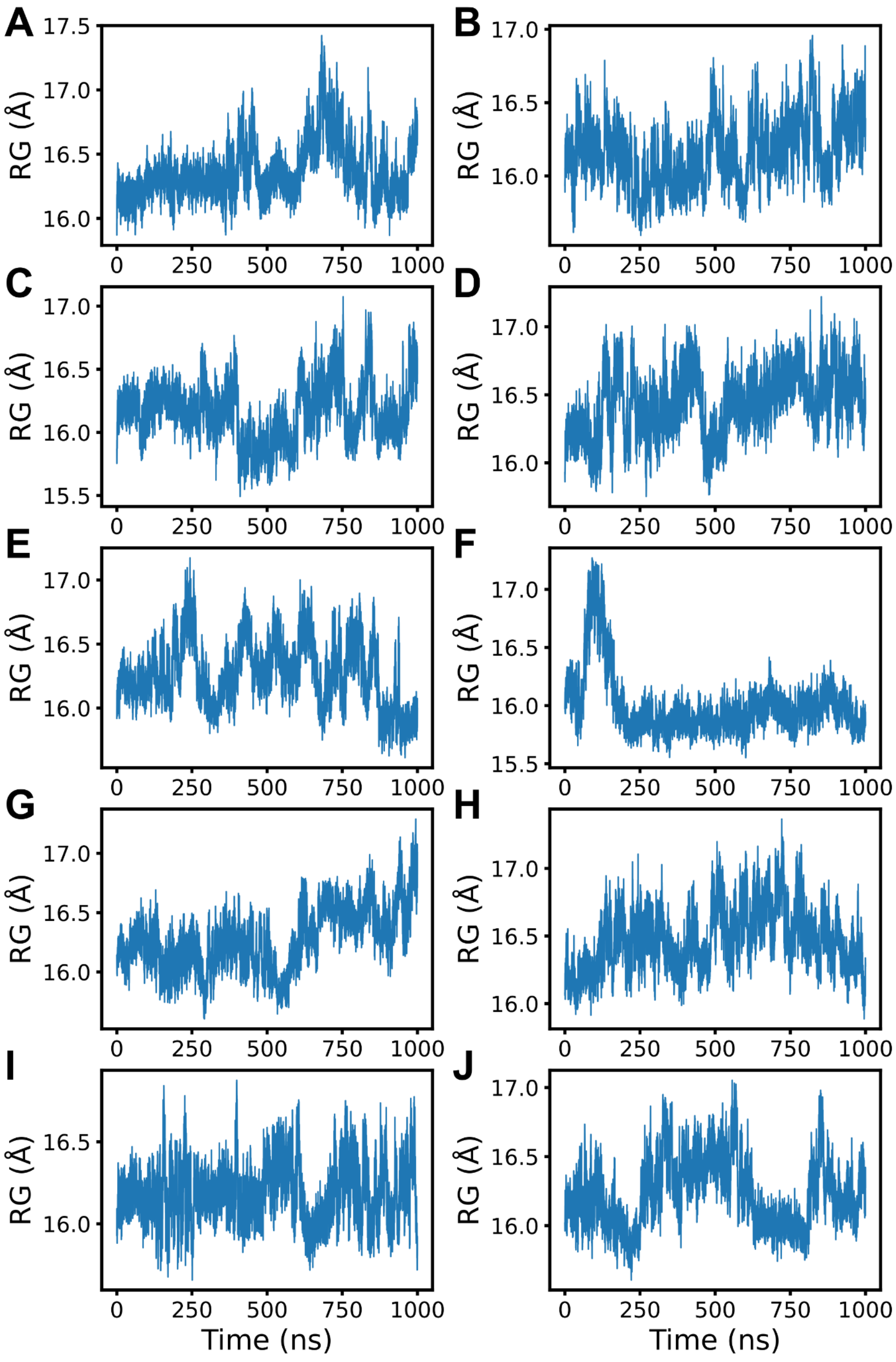


**Figure S 12 - Free TcmN molecular dynamics simulations.** Radius of gyration (Rg) of the MD trajectory for each replicate.

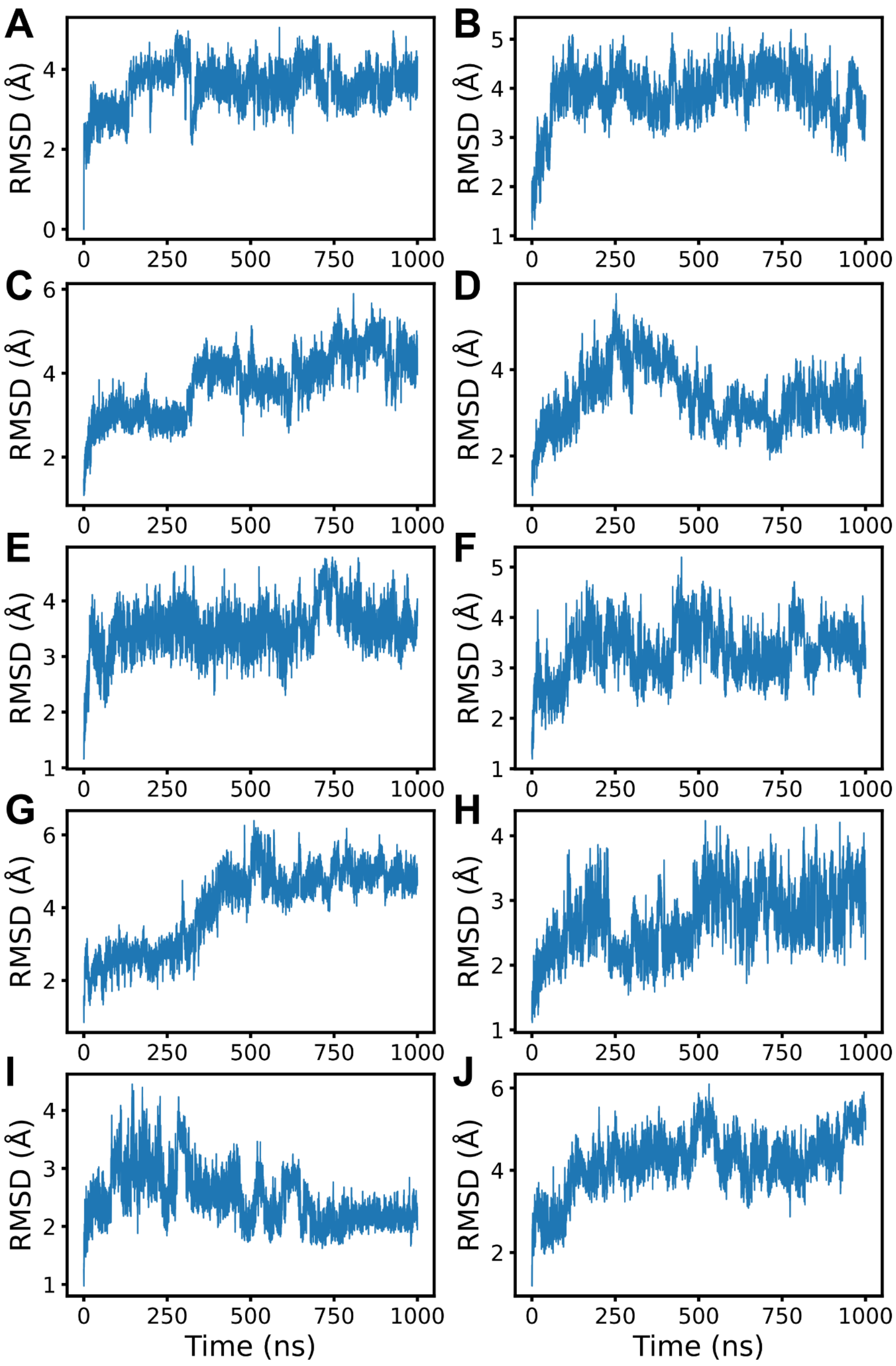


**Figure S 13 - INT12-bounded TcmN molecular dynamics simulations.** Root mean square deviation (RMSD) of the MD trajectory for each replicate.

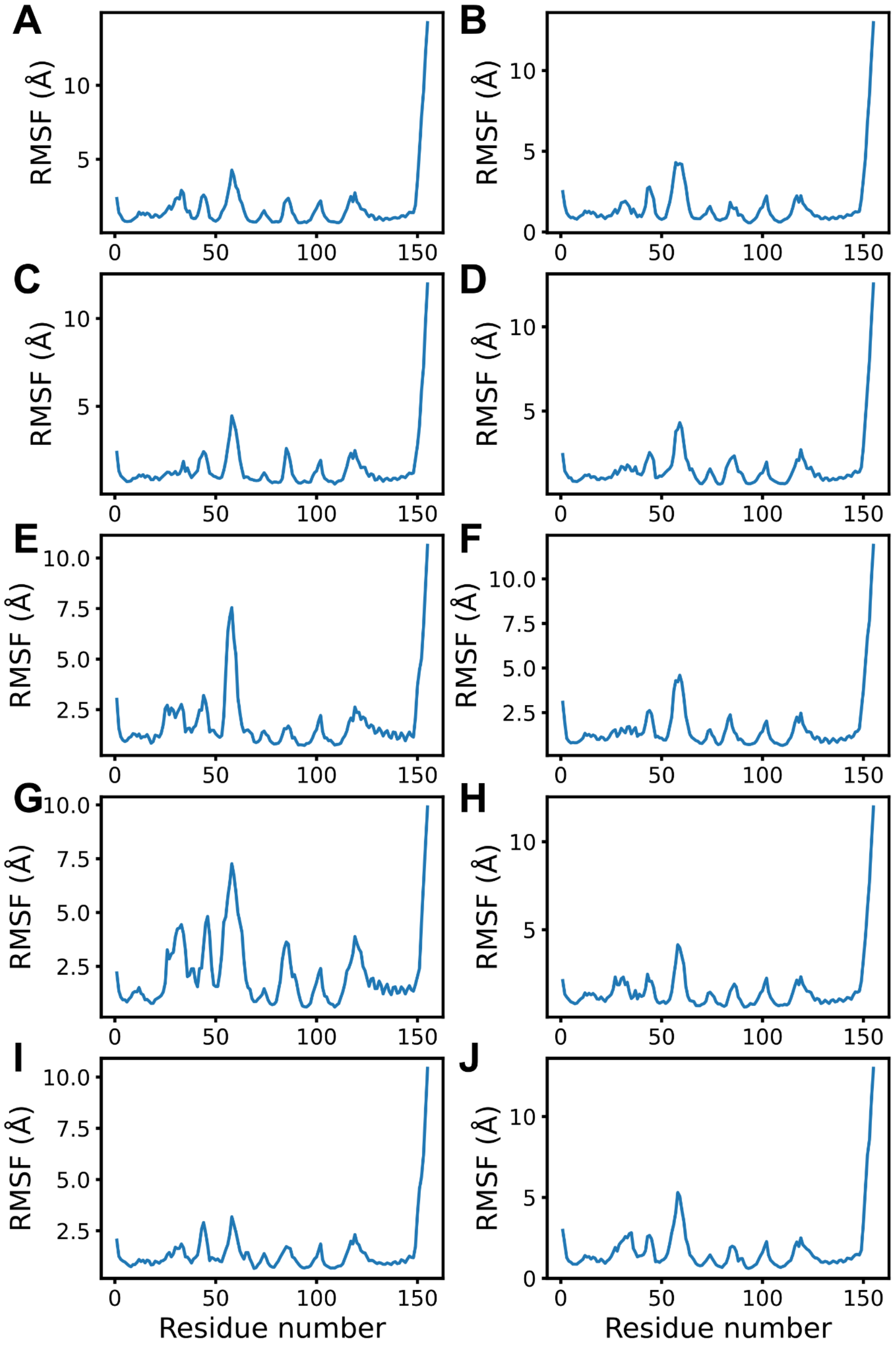


**Figure S 14 - TcmN with INT12 molecular dynamics simulations.** Root mean square fluctuation (RMSF) of the MD trajectory for each replicate.

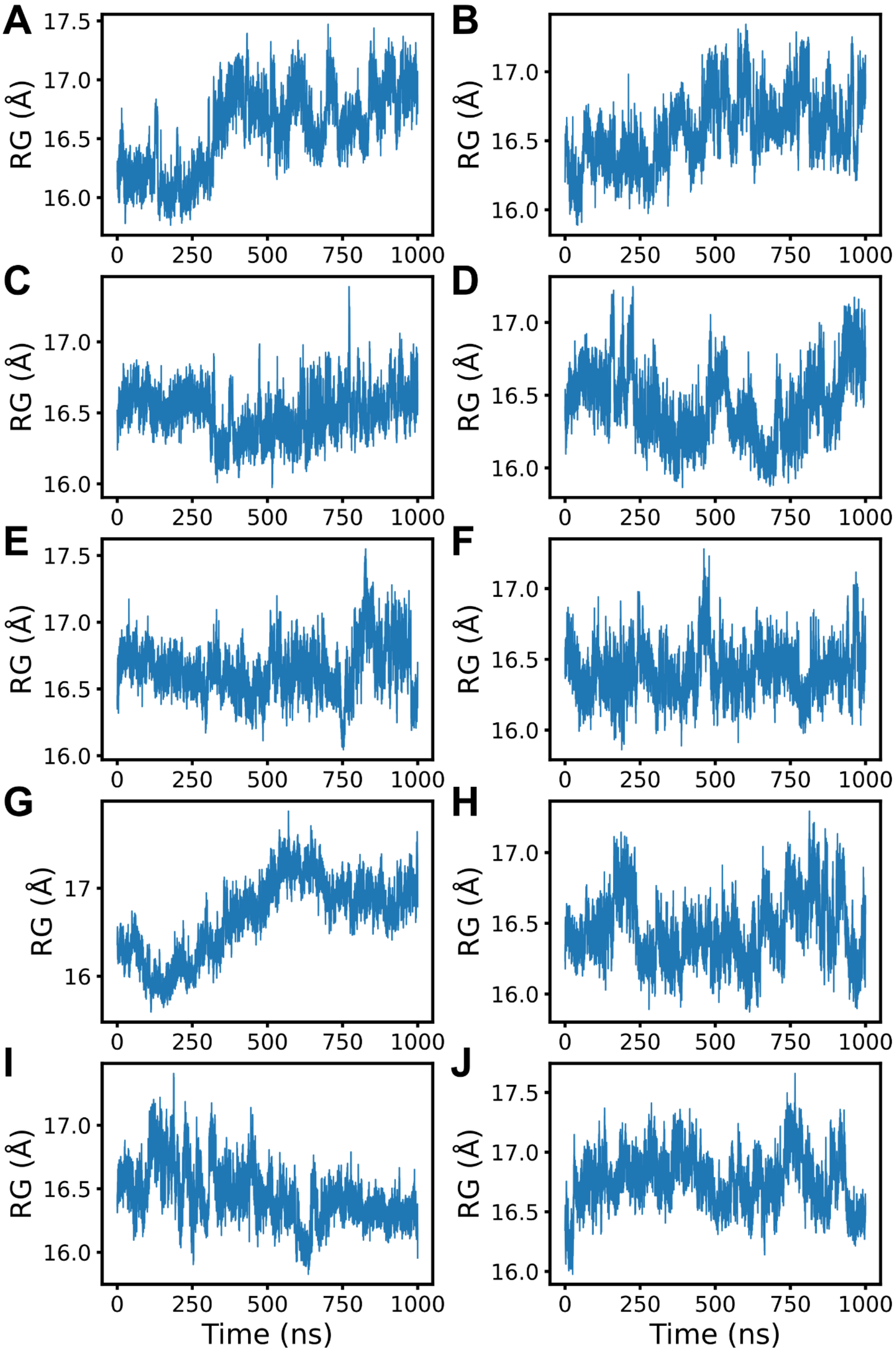


**Figure S 15 – INT12-bounded TcmN molecular dynamics simulations.** Radius of gyration (Rg) of the MD trajectory for each replicate.

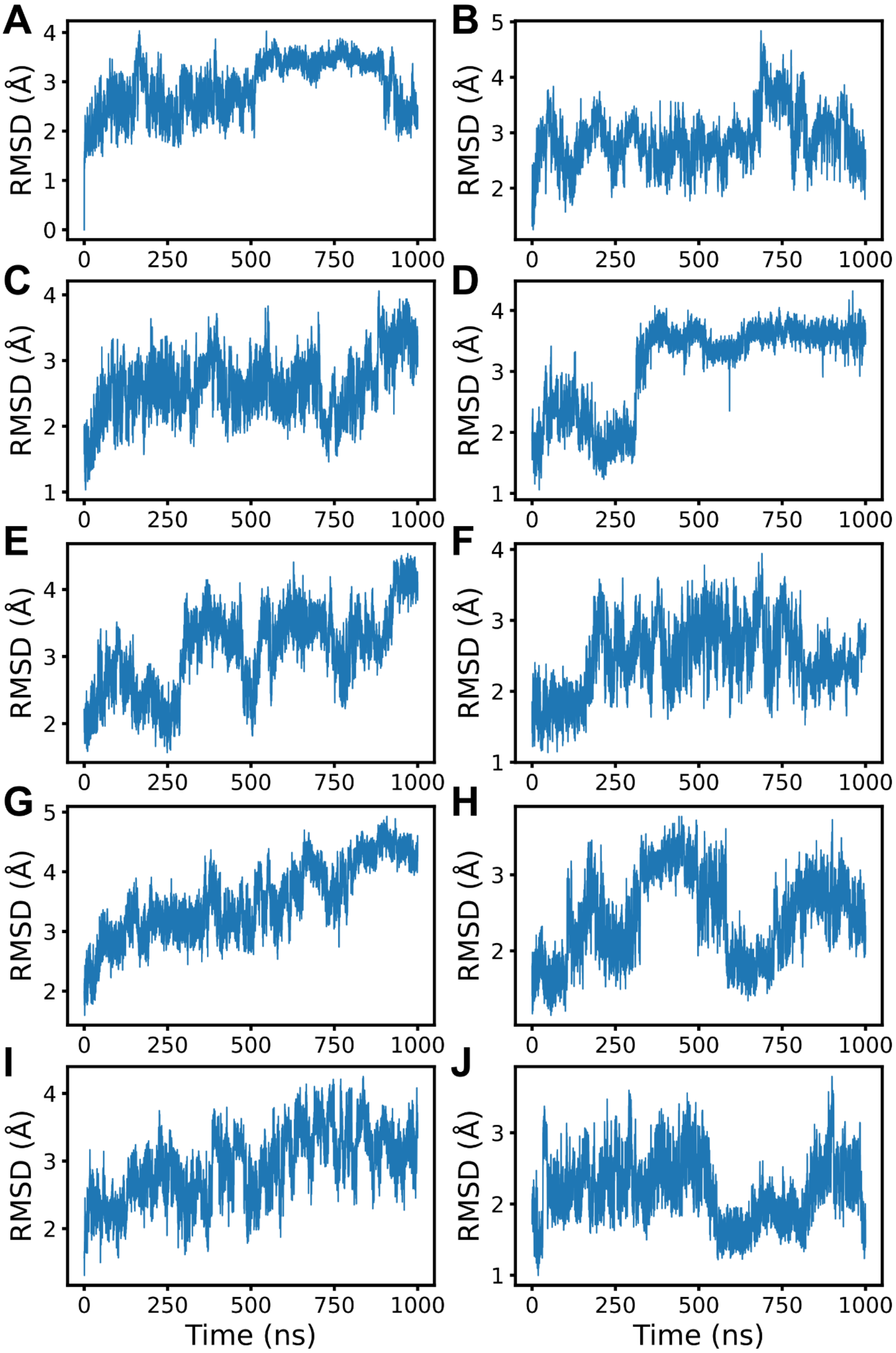


**Figure S 16 - (S)-Naringenin-bound TcmN molecular dynamics simulations.** Root mean square deviation (RMSD) of the MD trajectory for each replicate.

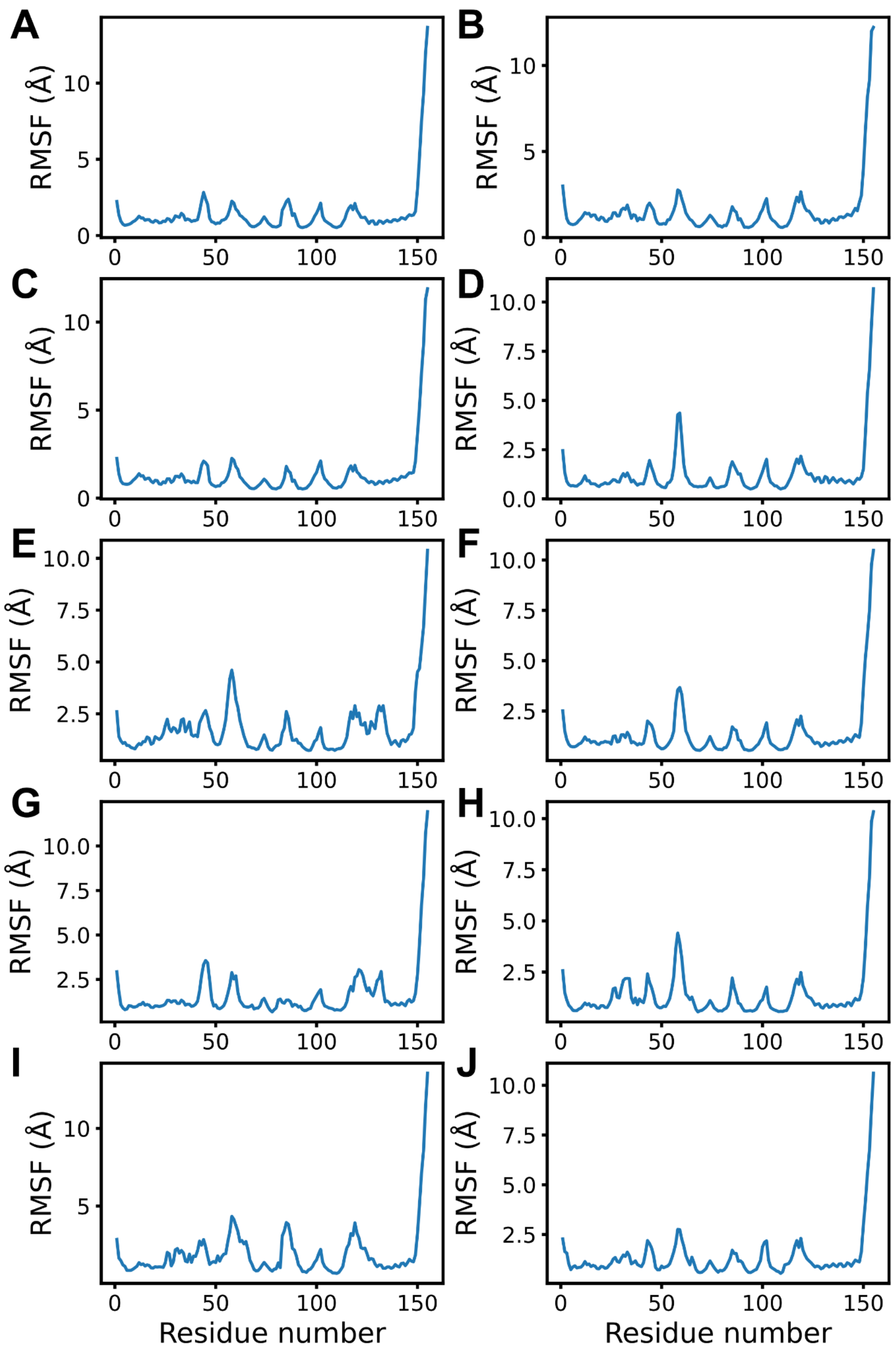


Figure S 17 – **(S)-Naringenin-bound TcmN** molecular dynamics simulations. Root mean square fluctuation (RMSF) of the MD trajectory for each replicate.

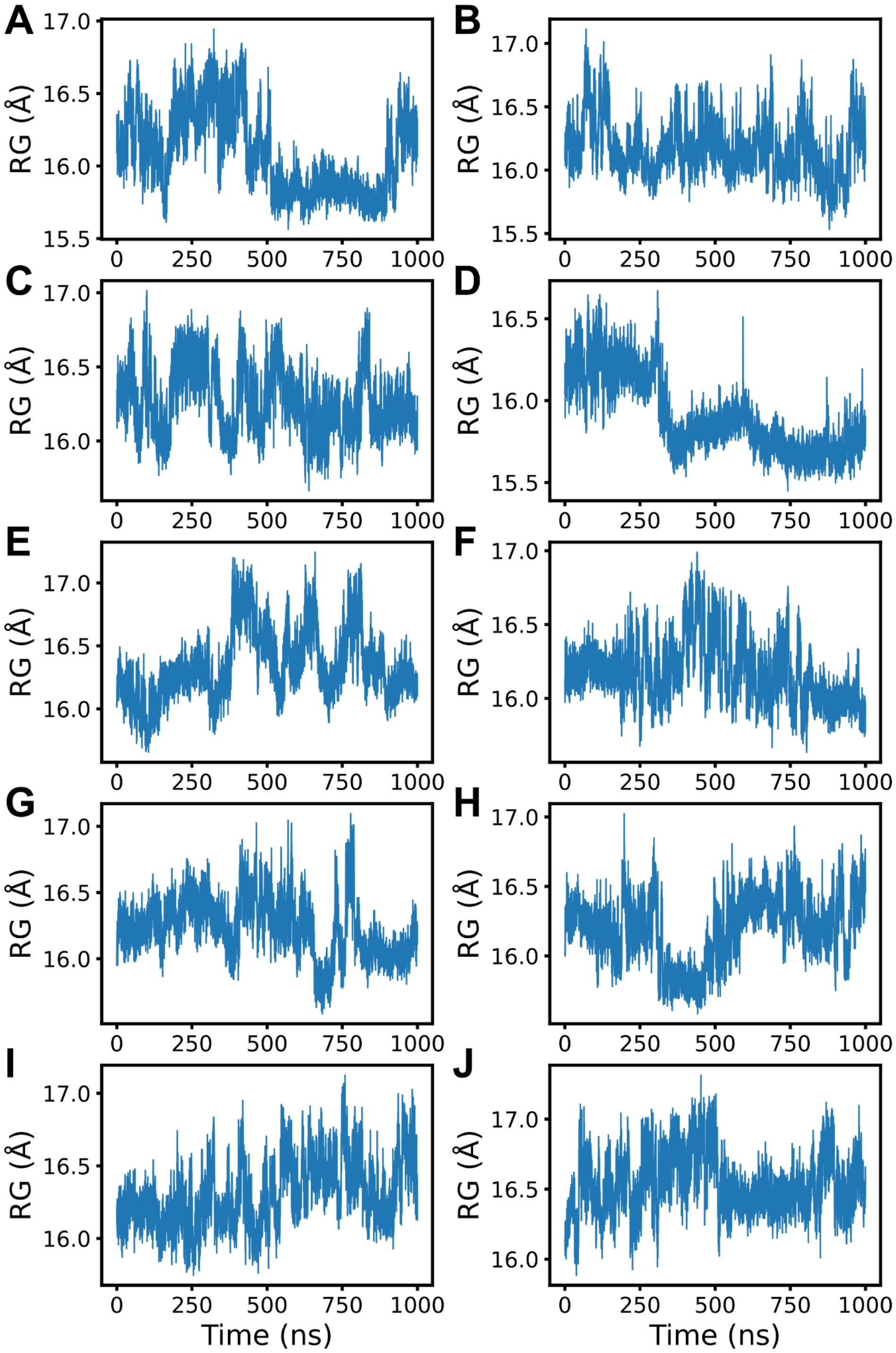


Figure S 18 **(S)-Naringenin-bound TcmN** molecular dynamics simulations. Radius of gyration (Rg) of the MD trajectory for each replicate.

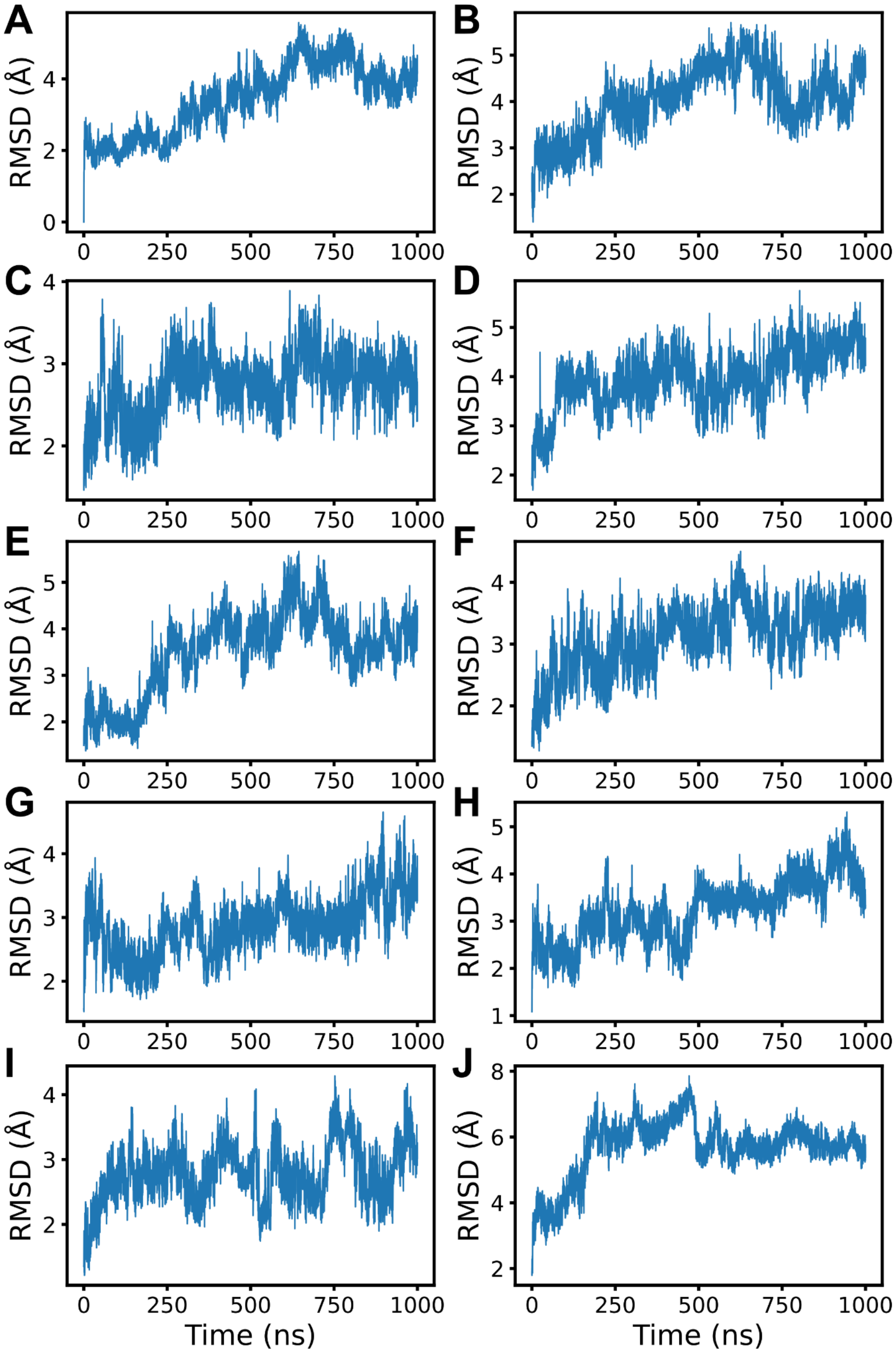


**Figure S 19 – TcmN bound to (R)-Naringenin (71w) molecular dynamics simulations.** Root mean square deviation (RMSD) of the MD trajectory for each replicate.

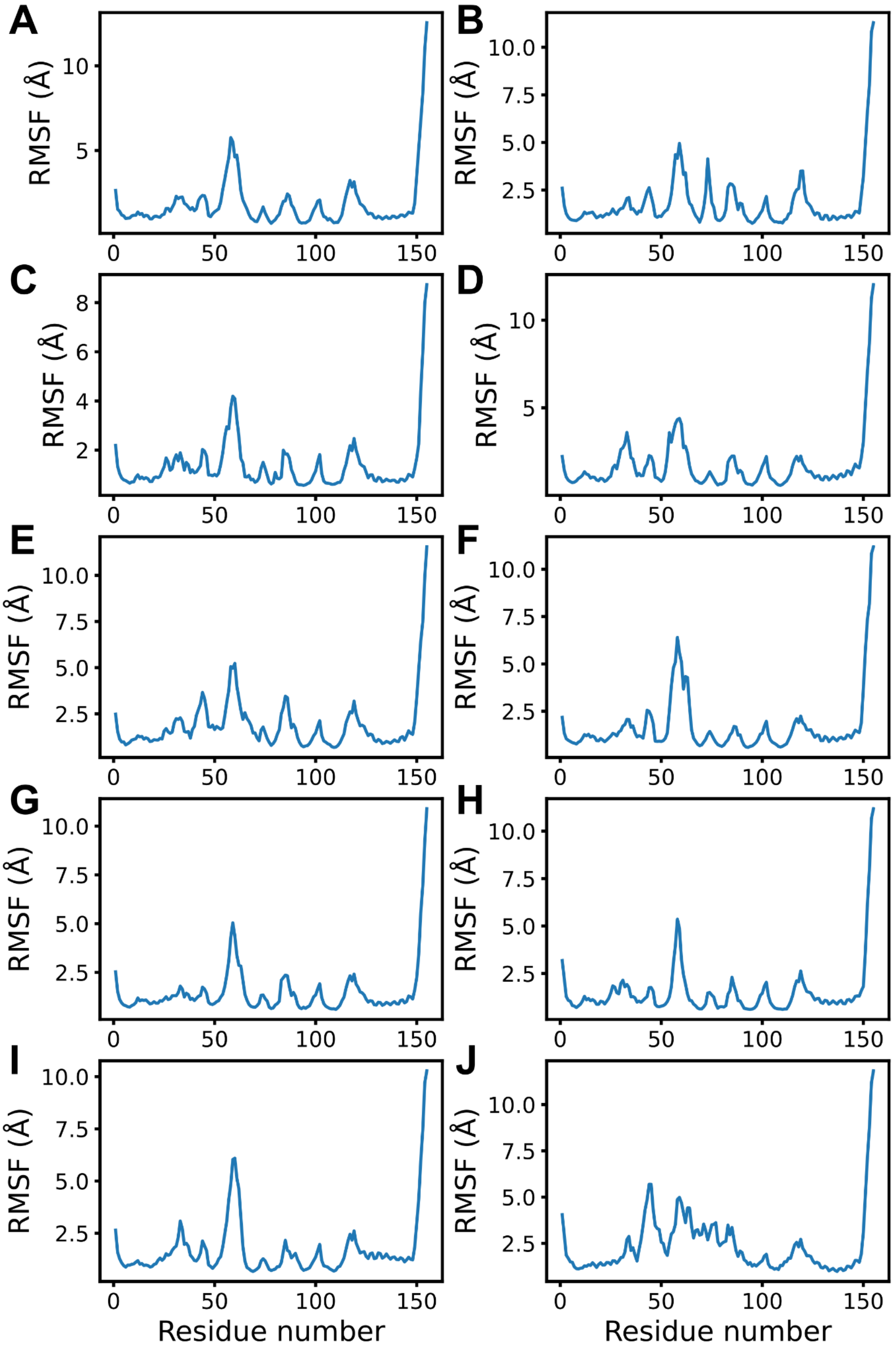


**Figure S 20 – TcmN bound to (R)-Naringenin (71w) molecular dynamics simulations.** Root mean square fluctuation (RMSF) of the MD trajectory for each replicate.

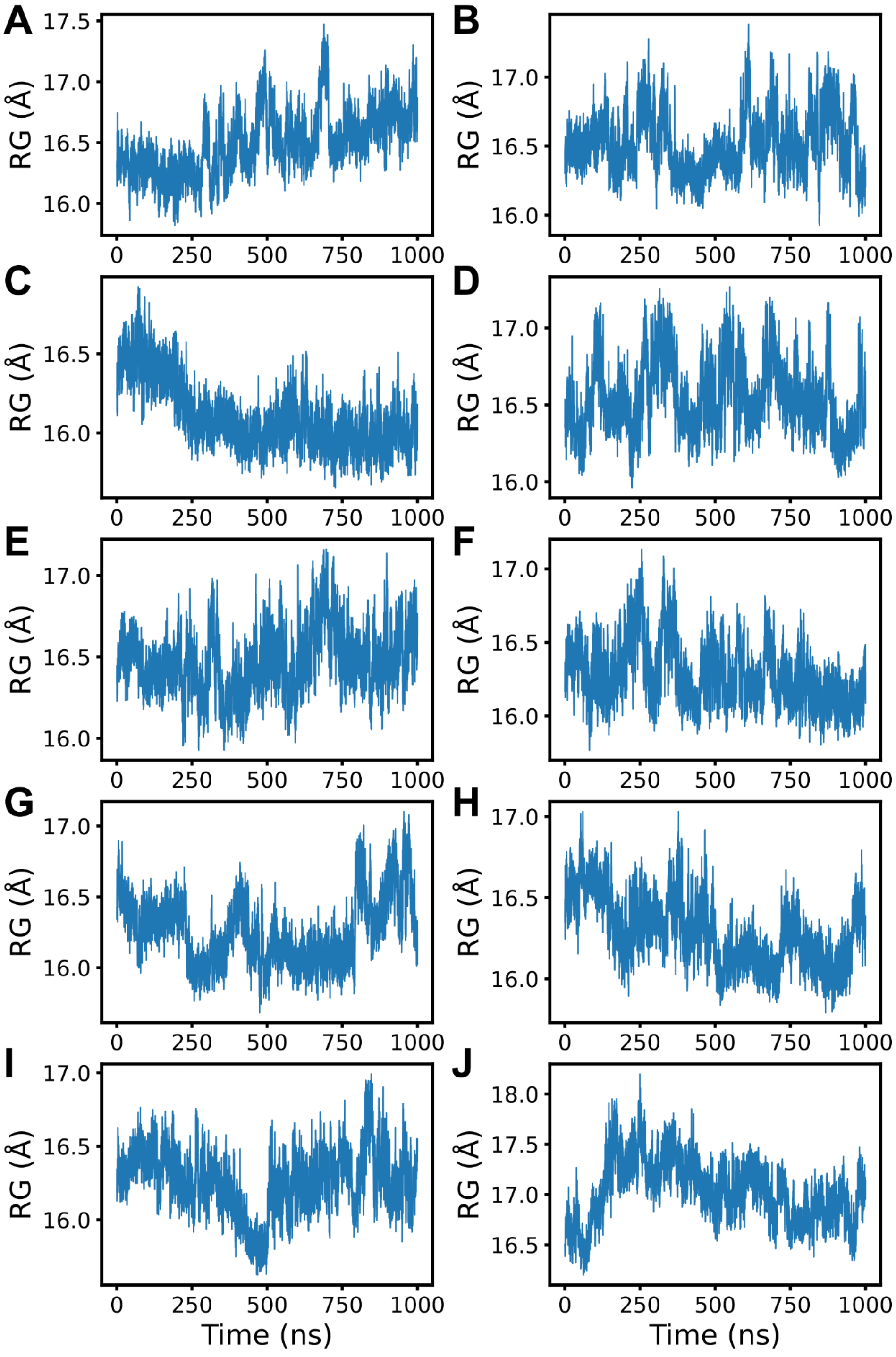


**Figure S 21 – TcmN bound to (R)-Naringenin (71w) molecular dynamics simulations.** Radius of gyration (Rg) of the MD trajectory for each replicate.

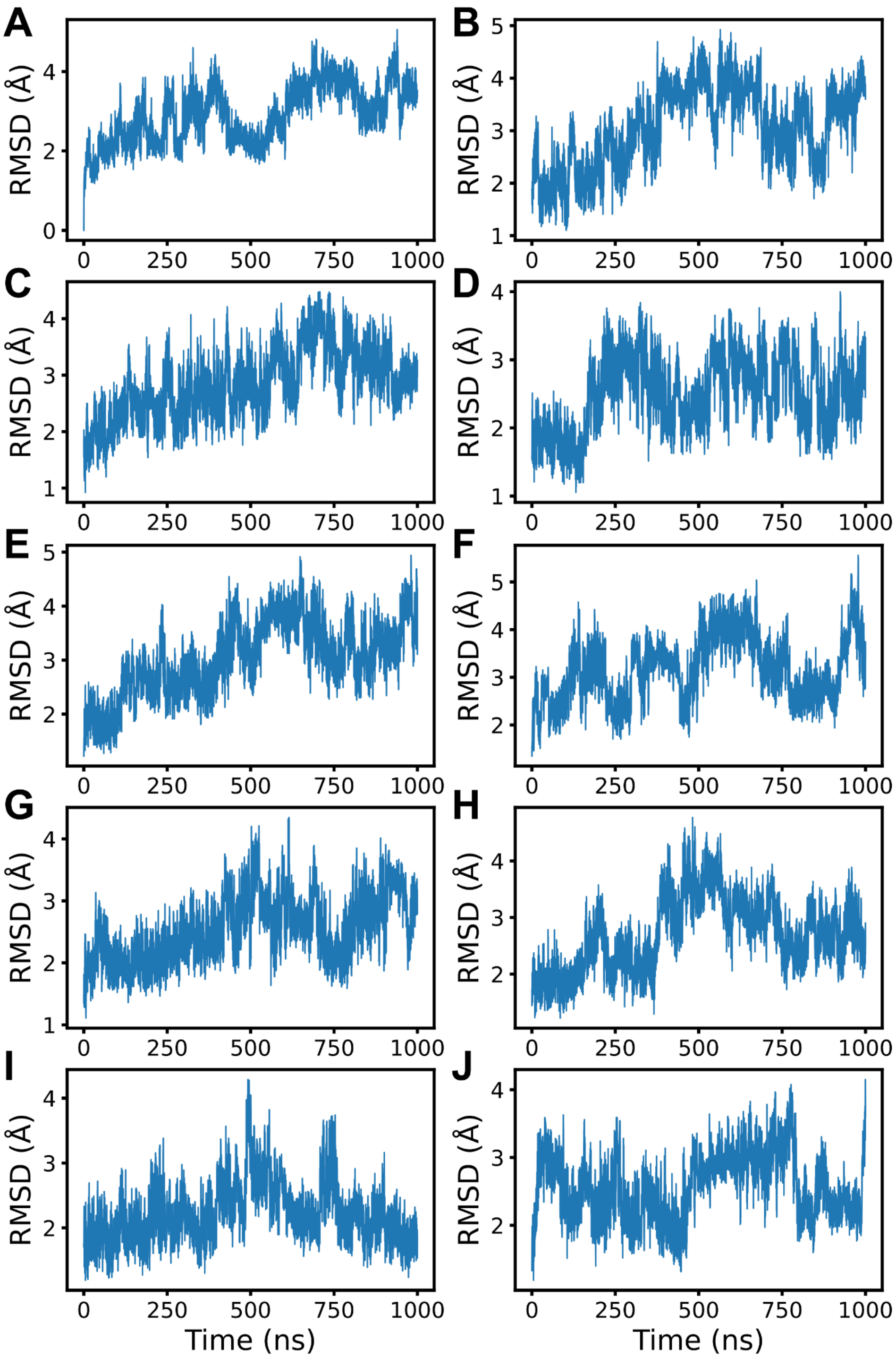


**Figure S 22 – TcmN bound to (R)-Naringenin (586w) molecular dynamics simulations.** Root mean square deviation (RMSD) of the MD trajectory for each replicate.

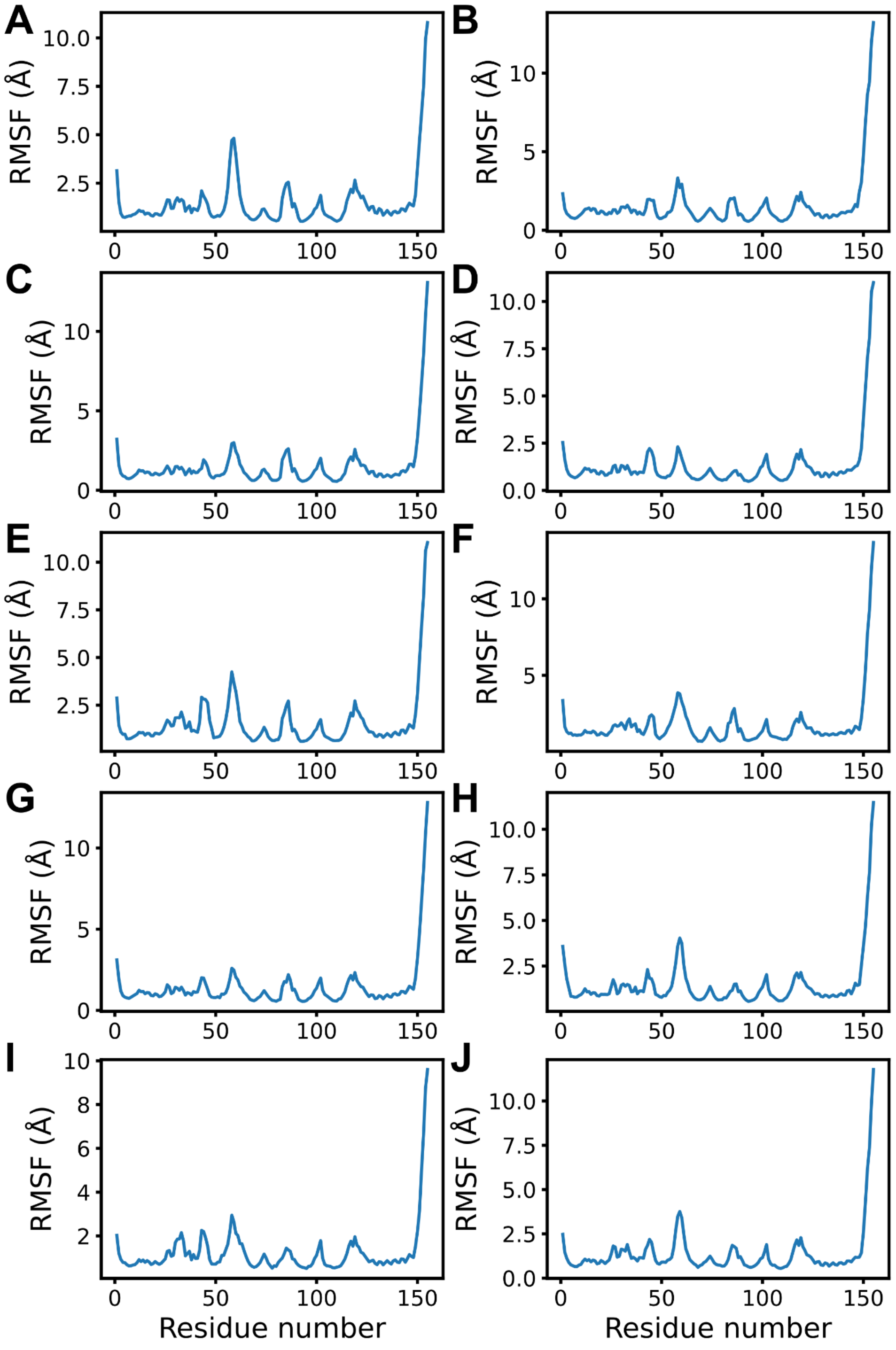


**Figure S 23 – TcmN bound to (R)-Naringenin (71w) molecular dynamics simulations.** Root mean square fluctuation (RMSF) of the MD trajectory for each replicate.

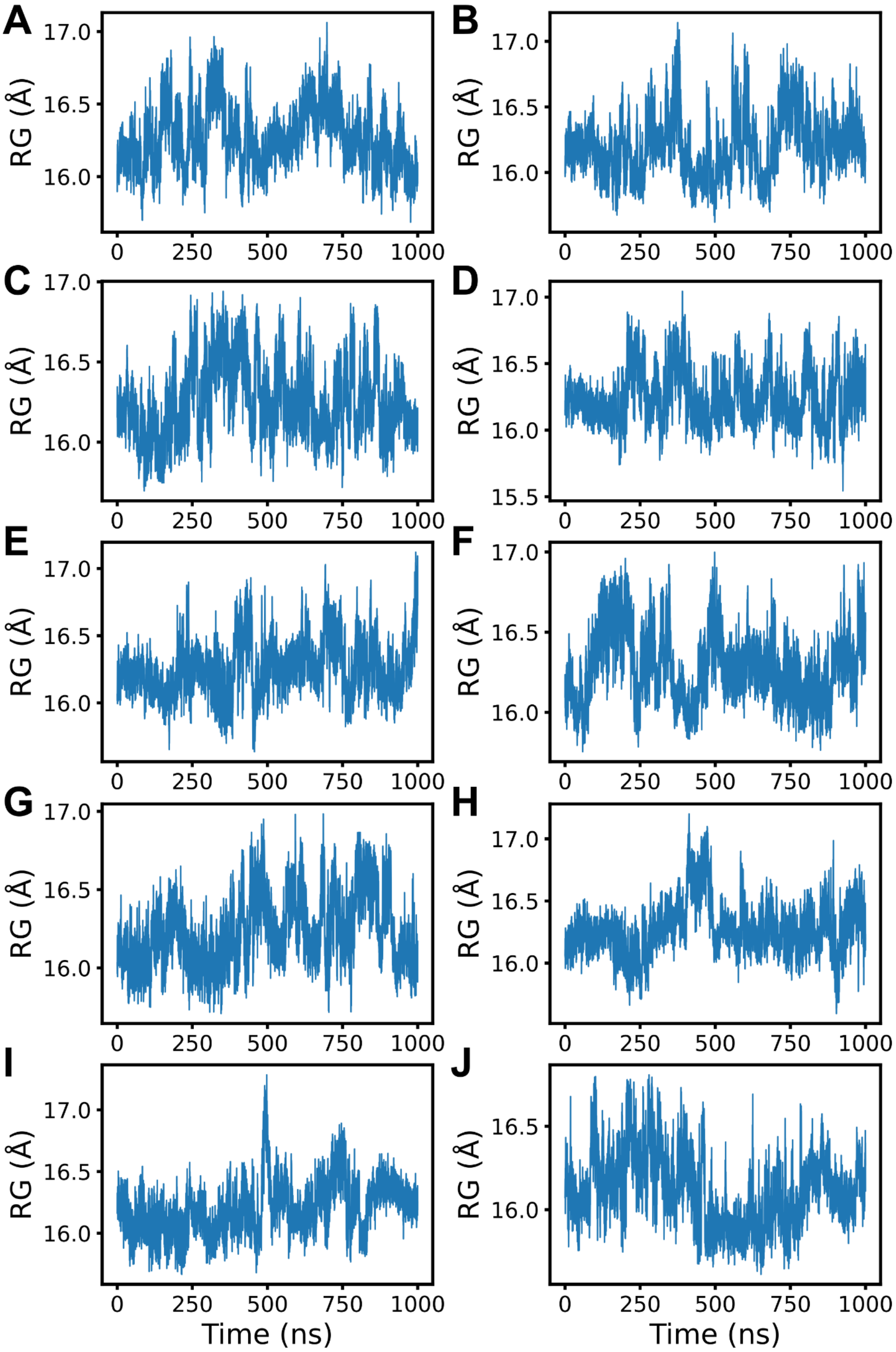


**Figure S 24 – TcmN bound to (R)-Naringenin (71w) molecular dynamics simulations.** Radius of gyration (Rg) of the MD trajectory for each replicate.

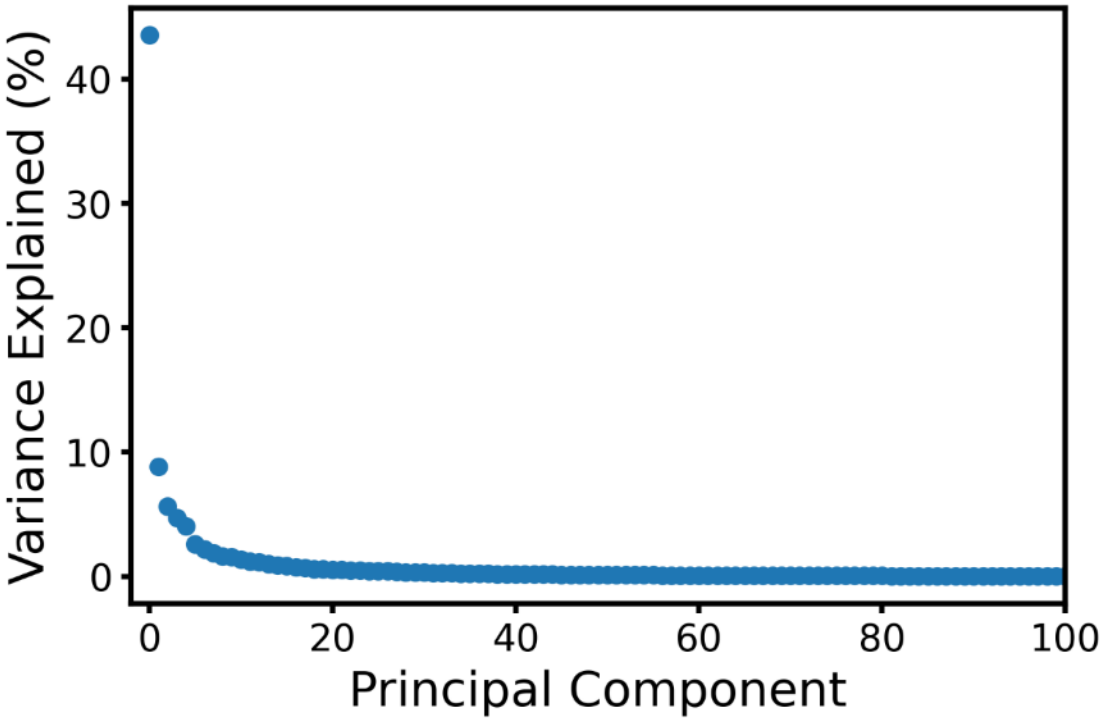


**Figure S 25 – Variance explained by each principal component (PC)**.

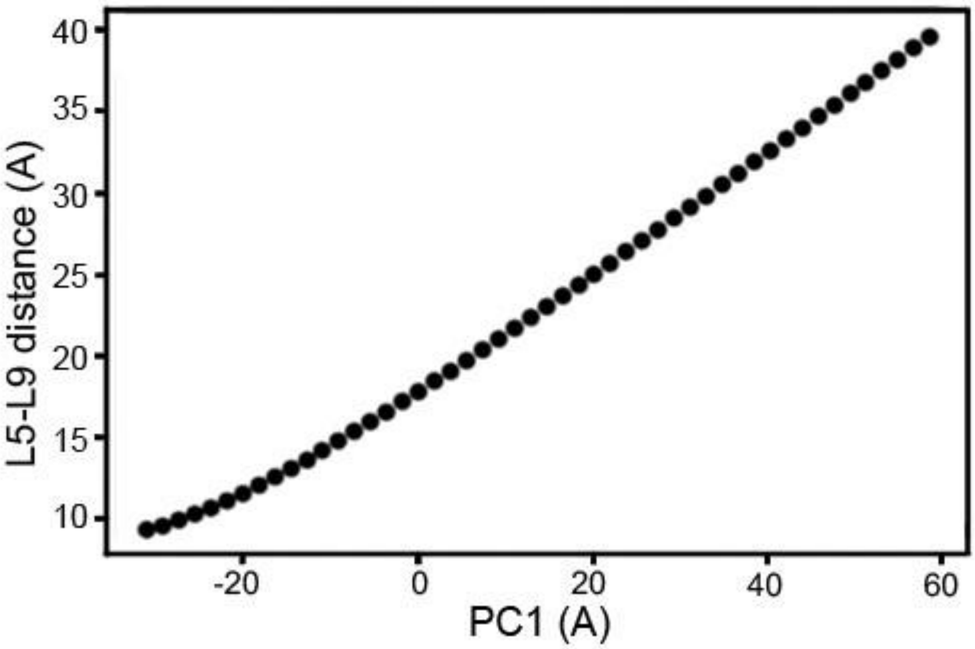


**Figure S 26 – PC1 correlation with L5-L9 distance**.

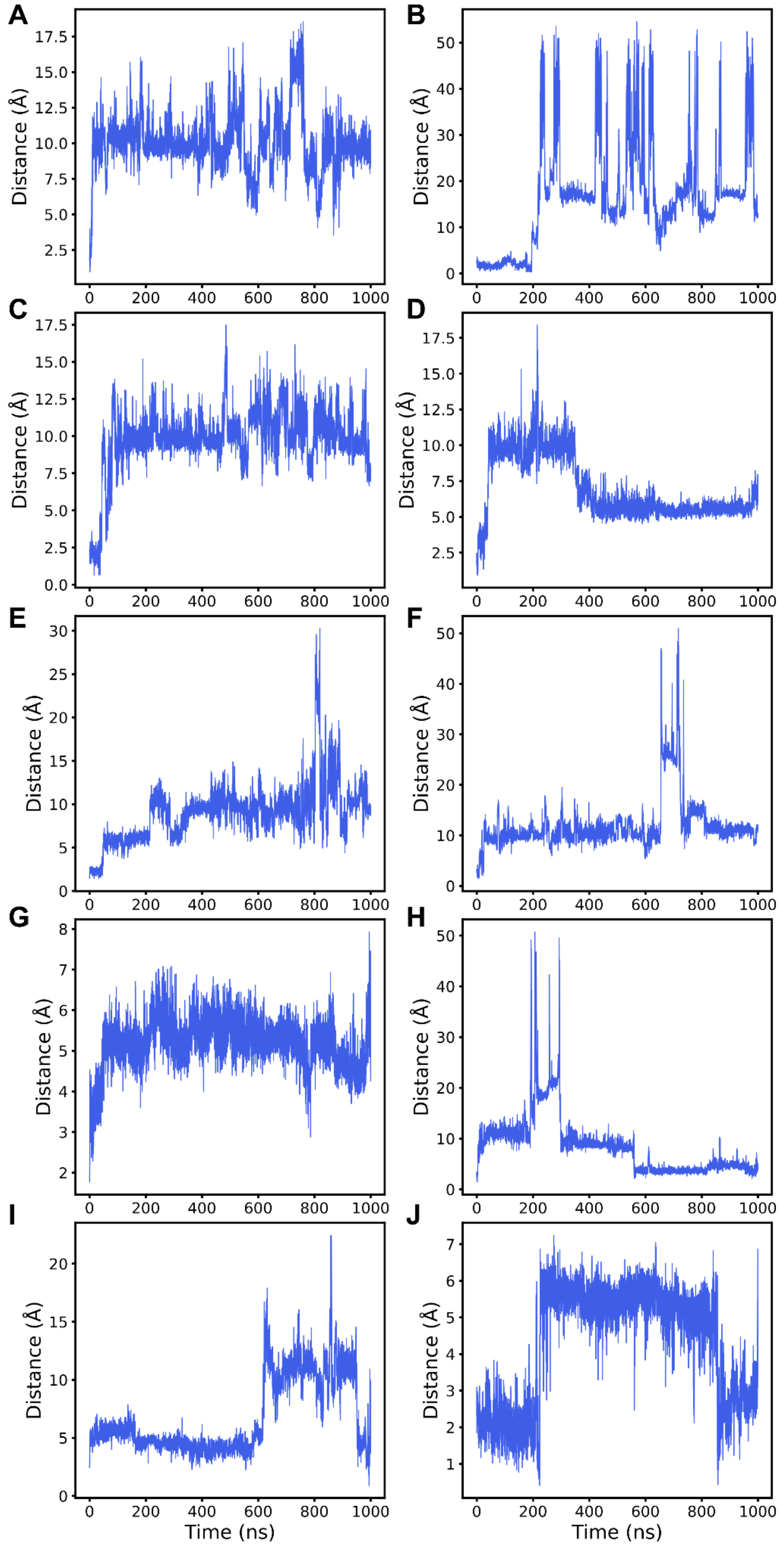


Figure S 27 – (S)-Naringenin distance from the TcmN cavity centroid in molecular dynamics simulations. Distances are shown for each replicate.

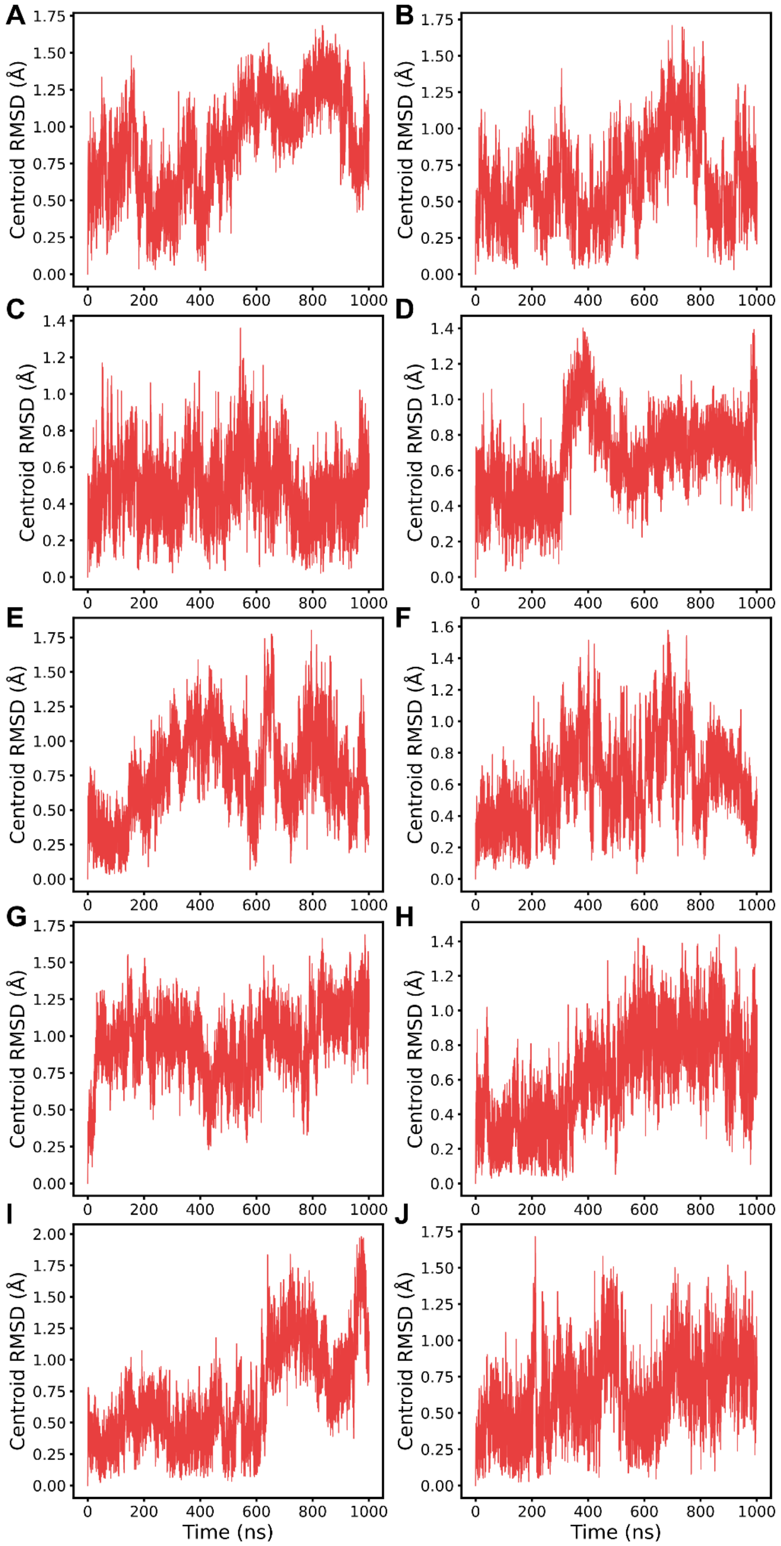


Figure S 28 – TcmN with bound (S)-Naringenin: cavity centroid RMSD in molecular dynamics simulations. Root mean square deviation (RMSD) of the cavity centroid is shown for each replicate.

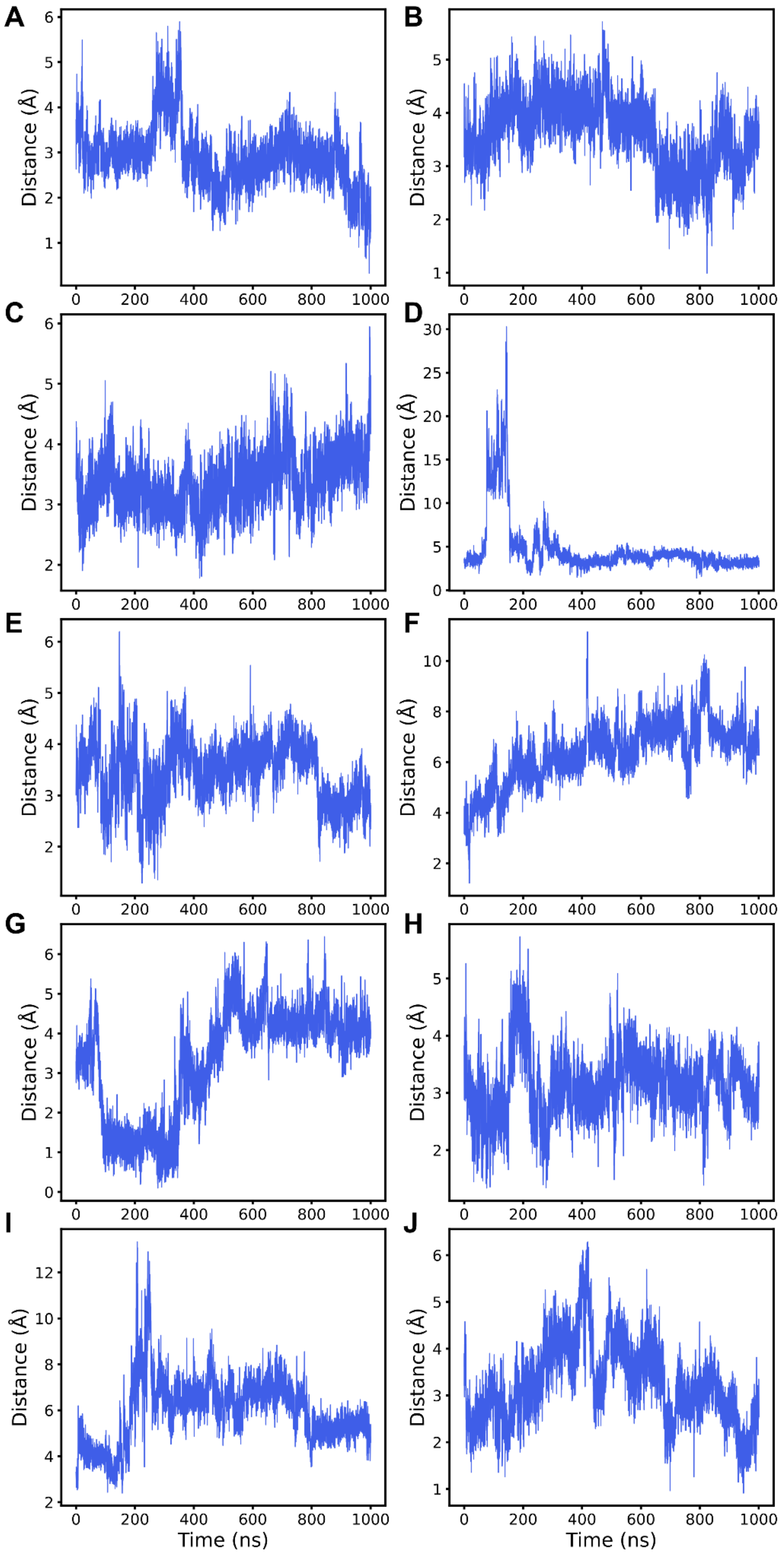


Figure S 29 – INT12 distance from the TcmN cavity centroid in molecular dynamics simulations. Distances are shown for each replicate.

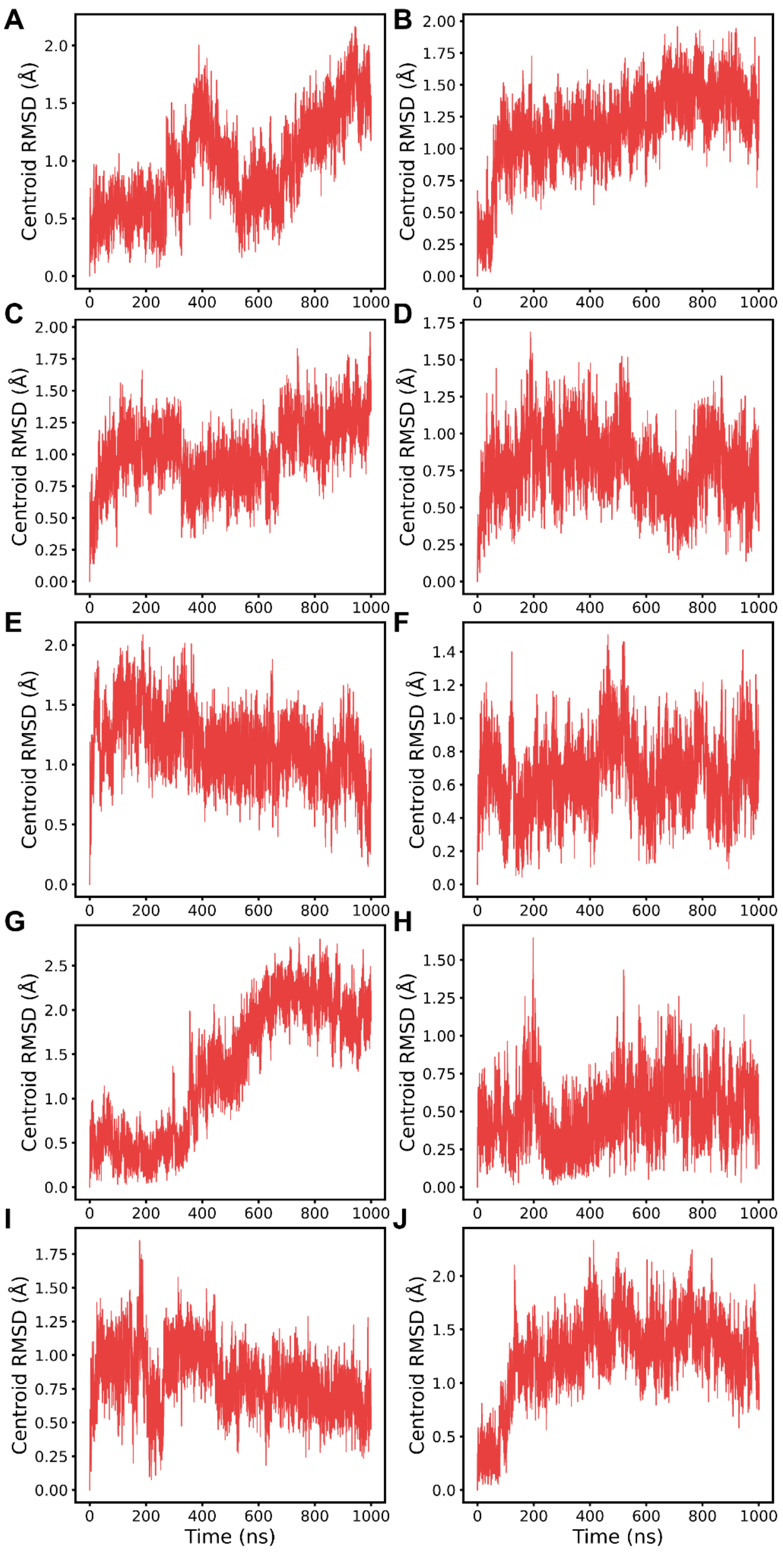


**Figure S 30 – TcmN with bound INT12: cavity centroid RMSD in molecular dynamics simulations.** Root mean square deviation (RMSD) of the cavity centroid is shown for each replicate.

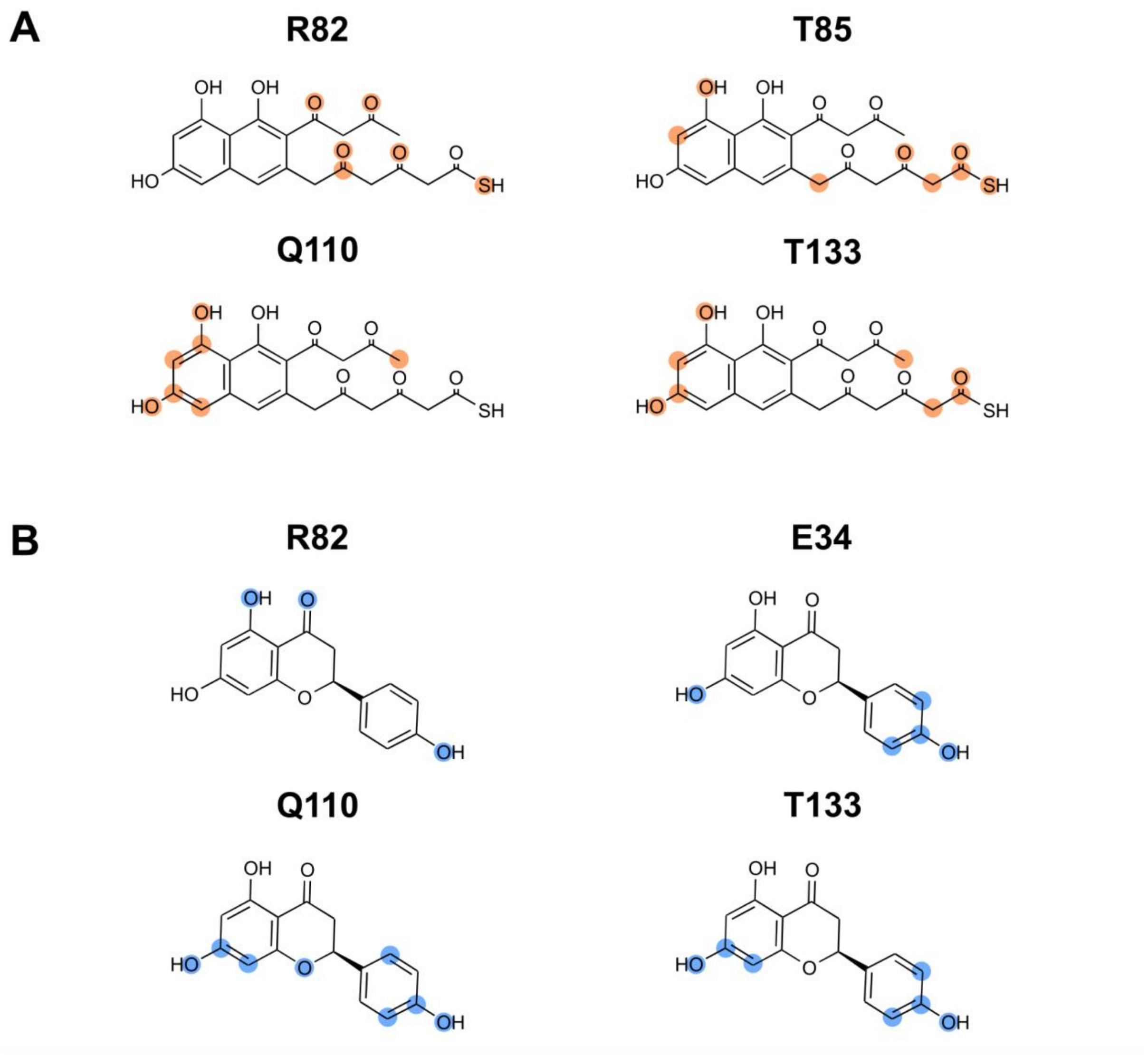


**Figure S 31 – Ligand heavy atoms forming the most frequent contacts with TcmN residues, identified from hydrogen bond occupancy.** (A) INT12. (B) (S)-Naringenin.

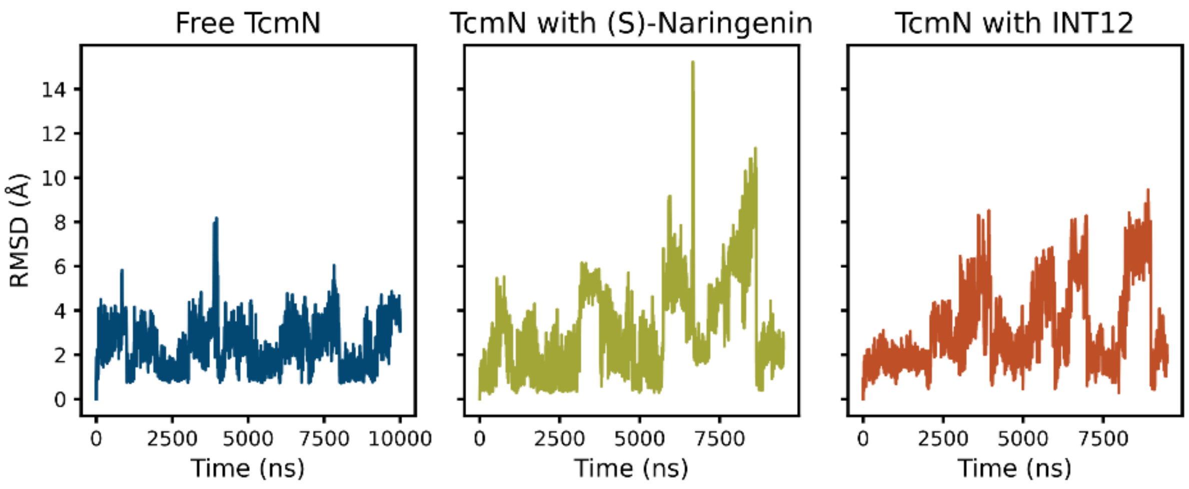


**Figure S 32 – Arg82 all heavy atom side chain RMSD.**
